# The selective autophagy receptor p62 and the heat shock protein HSP27 facilitate lysophagy via the formation of phase-separated condensates

**DOI:** 10.1101/2022.07.10.499468

**Authors:** Elizabeth R. Gallagher, Erika L.F. Holzbaur

## Abstract

Lysosomes are membrane-bound organelles that regulate cellular proteostasis. Loss of lysosomal integrity initiates cell death pathways. Thus, cells must rely on quality control mechanisms for protection, including the selective isolation and degradation of damaged lysosomes by lysophagy. Here, we report that the selective autophagy receptor SQSTM1/p62 is an essential lysophagy receptor recruited to damaged lysosomes in both HeLa cells and neurons. p62 oligomers form liquid-like condensates that are critical in lysophagy. These condensates are regulated by the small heat shock protein HSP27, which binds p62 to prevent p62 aggregation and facilitate autophagosome formation. Mutations in p62 are implicated in Amyotrophic Lateral Sclerosis (ALS), and expression of ALS-associated mutations in p62 impair lysophagy, suggesting that deficits in this pathway may contribute to the cellular pathogenesis of ALS. Thus, p62 oligomers cooperate with HSP27 to promote lysophagy by forming a platform for autophagosome biogenesis at damaged lysosomes.

## Introduction

Lysosomes are the primary degradative organelle in mammalian cells, responsible for the enzymatic digestion and recycling of macromolecules (Appelqvist et al., 2013). Lysosomal dysfunction and permeabilization endanger cellular health, risking release of calcium, degradative enzymes, and reactive oxygen species into the cytosol and initiating cell death pathways (Aits and Jaattela, 2013; Duve, 1983; Gabandé-Rodríguez et al., 2019; Repnik et al., 2014; Song et al., 2017).

Lysosomal dysfunction is particularly injurious to the central nervous system, as shown by the emerging link between disruption of lysosomal health and neurodegenerative disease (Malik et al., 2019; Monaco and Fraldi, 2020; Nixon, 2013; Usenovic and Krainc, 2012; Wallings et al., 2019). Disorders linked to lysosomal dysfunction include Lysosomal Storage Disorders, Parkinson’s Disease, Alzheimer’s Disease, Frontotemporal dementia, and Amytrophic Lateral Sclerosis (ALS) (Monaco and Fraldi, 2020; Root et al., 2021; Udayar et al., 2022). Lysosomal rupture has specifically been identified in Niemman-Pick Type A, Gaucher’s Disease, and Parkinson’s Disease (Freeman et al., 2013; Gabandé-Rodríguez et al., 2014; Jiang et al., 2017; Vitner et al., 2010). As lysosomal rupture can be extremely damaging to the cell, quality control mechanisms are engaged to rescue lysosomal integrity and protect the cell from lysosome-mediated cell death. However, there are few established mechanisms of neuronal lysosomal quality control (Eapen et al., 2021; Liu et al., 2020a).

Lysosomal quality control begins with an attempt to repair damaged lysosomes via the <u>E</u>ndosomal <u>S</u>orting <u>C</u>omplexes <u>R</u>equired for <u>T</u>ransport (ESCRT) machinery (Radulovic et al., 2018; Skowyra et al., 2018). In the absence of lysosomal repair, ruptured lysosomes are targeted for degradation via selective autophagy, referred to as lysophagy (Jia et al., 2020a; Radulovic et al., 2018; Skowyra et al., 2018). Selective autophagy is a process in which autophagosomes, double-membraned vesicles, form in the cytoplasm to sequester and ultimately degrade cellular cargo (Chen et al., 2019; Khaminets et al., 2016; Stolz et al., 2014).

Damaged lysosomes undergo either ubiquitin-dependent or ubiquitin-independent lysophagy (Chauhan et al., 2016; Di Rienzo et al., 2020; Maejima et al., 2013; Papadopoulos and Meyer, 2017; Papadopoulos et al., 2017; Radulovic et al., 2018). In ubiquitin-dependent lysophagy, the ubiquitination of lysosomal proteins drives the recruitment of selective autophagy receptors (Maejima et al., 2013; Papadopoulos et al., 2017; Radulovic et al., 2018). These receptors link damaged lysosomes to the growing autophagosome and facilitate their selective engulfment (Chen et al., 2019; Stolz et al., 2014). However, there have been conflicting reports regarding which receptors are involved in lysophagy (Eapen et al., 2021; Maejima et al., 2013; Papadopoulos et al., 2017).

Several selective autophagy receptors have been identified on the surface of damaged endolysosomes, including NDP52, TAX1BP1, and p62/SQSTM1 (Eapen et al., 2021; Fujita et al., 2013; Hung et al., 2013; Koerver et al., 2019; Maejima et al., 2013; Yoshida et al., 2017). The receptor p62 is of particular interest, given it has been shown to be essential in the selective autophagy of protein aggregates (aggrephagy) (Stolz et al., 2014), but not for the clearance of dysfunction mitochondria via mitophagy, despite localizing to dysfunctional mitochondria (Narendra et al., 2010). While p62 is recruited to damaged lysosomes (Maejima et al., 2013), evidence is lacking on whether p62 is essential to promote lysophagy. This question is of interest, because a major, autophagy-promoting role of p62 in aggrephagy is the formation of liquid-liquid phase separated condensates (Sun et al., 2018; Turco et al., 2019). However, the significance of p62 condensates in organellophagy remains unknown. p62 is not required for Parkin-mediated mitophagy, although p62 drives the clustering of dysfunctional mitochondria (Narendra et al., 2010; Wong and Holzbaur, 2014). Conversely, in Parkin-independent mitophagy, p62 condensates appear critical (Peng et al., 2021). These data suggest that p62 condensates are unique to some forms of selective autophagy, prompting the question: are p62 condensates required in lysophagy?

Here, we demonstrate that p62 functions as an essential lysophagy receptor, responding to lysosomal damage in HeLa cells as well as human iPSC-derived neurons and rat hippocampal neurons. This response is selective, which we demonstrate using the genetically encoded lysosomal photosensitizer KillerRed to focally damage lysosomes. We find that lysophagy requires p62 oligomerization, as loss of p62 self-association prevents its recruitment to damaged lysosomes and impairs engulfment of the organelles by mAtg8-positive autophagosomes. Maintaining the liquid phase properties of p62 is critical; we find that the small heat shock protein HSP27 responds to lysosomal damage downstream of p62, maintains the liquid-like state of p62 oligomers by limiting p62 aggregation, and thus promotes autophagosome formation. ALS-associated mutations in p62 disrupt lysophagy in cellular assays, further implicating defects in lysosomal quality control in neurodegenerative disease. Thus, we propose that p62 facilitates lysophagy via condensate formation that is regulated by HSP27, forming a platform for *de novo* autophagosome biogenesis to rapidly and effectively engulf damaged lysosomes.

## Results

### p62 is dynamically recruited to damaged lysosomes in HeLa cells

To address the role of p62 in lysophagy, we used the well-characterized lysomotropic agent L-Leucyl-L-Leucine methyl ester (LLOMe) to induce acute lysosomal damage and used immunofluorescence to assess p62 recruitment (Jia et al., 2018, 2020b, 2020a; Papadopoulos et al., 2017; Radulovic et al., 2018; Skowyra et al., 2018; Thiele and Lipsky, 1990; Villamil Giraldo et al., 2014). We treated HeLa cells with either ethanol (EtOH) as a vehicle control or LLOMe at either 250µM or 750µM LLOMe for 2hrs. We then evaluated the recruitment of endogenous p62 to lysosomes, identified as organelles positive for the lysosomal transmembrane protein LAMP1 (Figure 1A) (Fukuda et al., 1988). In accordance with previous reports, we observed significant increases in the fraction of lysosomal area occupied by endogenous p62 when treated with 250µM or 750µM LLOMe (EtOH: 1.1 ± 0.43%; 250µM LLOMe: 20 ± 2.3%; 750µM LLOMe: 25 ± 0.68%) (Figure 1A,B), with no difference in average cellular area of LAMP1 among the conditions (Figure 1C) (Koerver et al., 2019; Papadopoulos et al., 2017).

**Figure 1.**
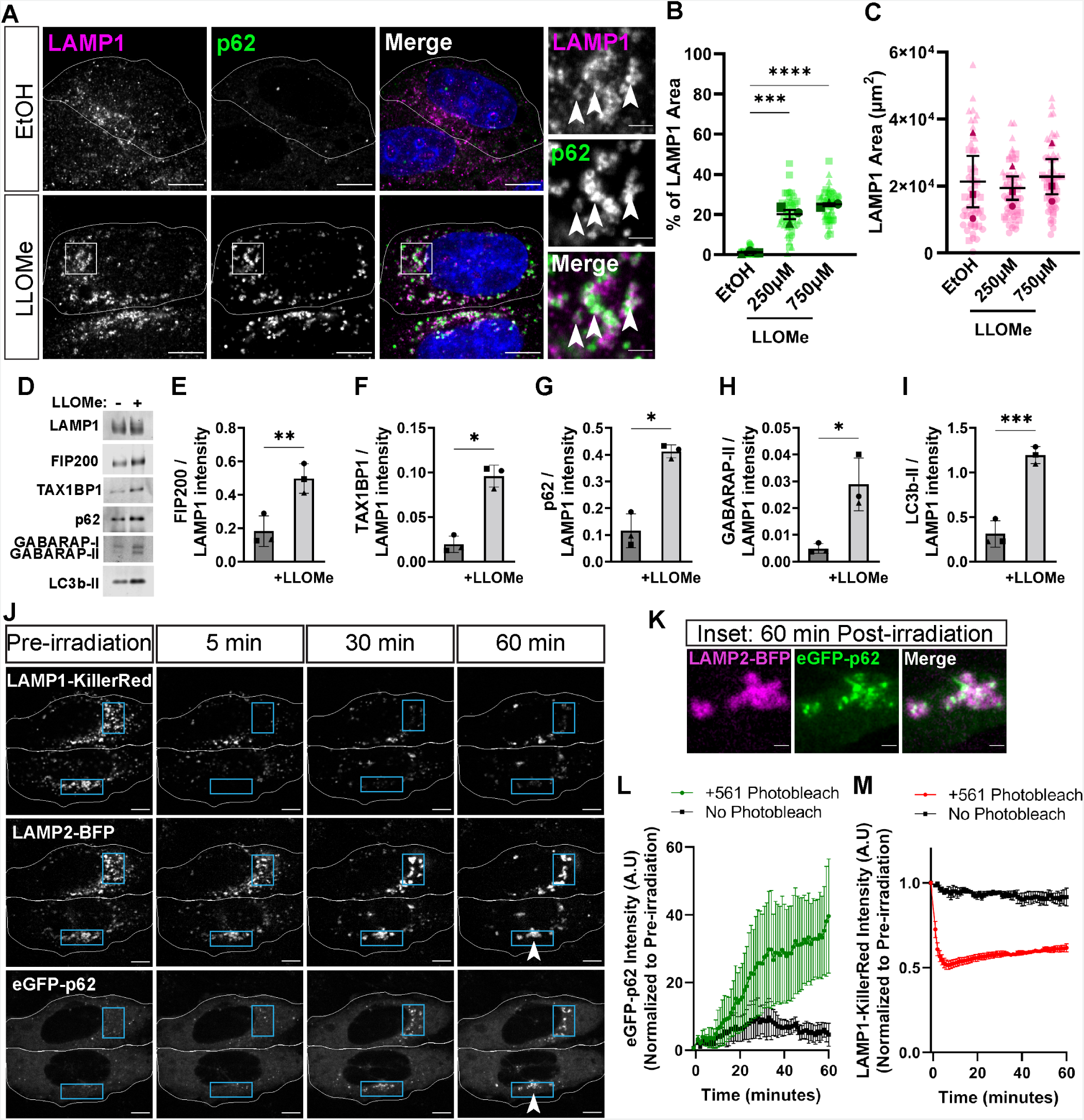
p62 is dynamically recruited to damaged lysosomes in HeLa cells. (A) Single z-plane representative images of anti-LAMP1 and anti-p62 in HeLa cells treated with ethanol (EtOH) as a vehicle control or LLOMe (at 250µM or 750 µM) for 2 hours. Cell outlines are traced. White boxes indicate inset image region. Arrowheads indicate regions of co-localization between LAMP1 and p62. Scale Bar: 10µm; inset: 2µm. (B) Quantification of the overlapping area from max projections between p62 and LAMP1 in the different conditions (One-way ANOVA, p_(EtOH/250)_ < 0.001, p_(EtOH/750)_ < 0.0001, N=3, n_(EtOH)_= 54, n_(250µM LLOMe)_=56, n_(750µM LLOMe)_=57). (C) Quantification of LAMP1 area from max projections in the different conditions (One-way ANOVA, p_(EtOH/250)_ = 0.96, p_(EtOH/750)_ = 0.9766, N=3, n_(EtOH)_= 54, n_(250µM LLOMe)_=56, n_(750µM LLOMe)_=57). (D) Western blot analysis of lysosomal immunoprecipitation following either no treatment or treatment with 1.0mM LLOMe for 1 hour. (E-I) Quantification of Lyso-IP. (E) Quantification of anti-FIP200 band intensities following normalization to anti-LAMP1 intensity (Unpaired t-test, p < 0.05, N=3). (F) Quantification of anti-TAX1BP1 band intensities following normalization to anti-LAMP1 intensity (Unpaired t-test, p < 0.01, N=3). (G) Quantification of anti-p62 band intensities following normalization to anti-LAMP1 intensity (Unpaired t-test, p < 0.05, N=3). (H) Quantification of anti-GABARAP band intensities following normalization to anti-LAMP1 intensity (Unpaired t-test, p < 0.05, N=3). (I) Quantification of anti-LC3b band intensities following normalization to anti-LAMP1 intensity (Unpaired t-test, p < 0.001, N=3). (J) Representative images of HeLa transiently transfected with LAMP1-KillerRed, LAMP2-BFP, and eGFP-p62. Images depict pre-and post-561nm laser irradiation. Cell outlines are traced. Blue boxes reflect regions irradiated. Arrowheads reflect location of inset images at 60min post-laser activation in Figure 1K. Scale Bar 10µm. (K) Inset images from Figure 1J. HeLa at 60min post-laser irradiation. Scale Bar: 2µm. (J) Quantification of eGFP-p62 intensities over one hour within regions either activated by laser irradiation or untreated. (K) Quantification of LAMP1-KillerRed intensities over one hour within regions either activated by laser irradiation or untreated.

As a parallel approach, we evaluated the association of p62 with damaged lysosomes using lysosomal immunoprecipitation following treatment with LLOMe (Abu-Remaileh et al., 2017). We generated a HeLa cell line stably expressing the lysosomal transmembrane protein TMEM192 C-terminally tagged with 3xHA. Cells were either untreated or treated with 1mM LLOMe for 1hr. We immunoprecipitated lysosomes and observed significant increases in lysosome-associated FIP200, TAX1BP1, p62, and lipidated Atg8-family proteins (GABARAP and LC3b) in LLOMe-treated cells (Figure 1D-I). These data further demonstrate that the autophagy receptors p62 and TAX1BP1 accumulate on damaged lysosomes within one hour, along with components of the core autophagy machinery.

Next, we examined the temporal dynamics of p62 recruitment following spatially-restricted lysosomal damage. We engineered a genetically encoded, lysosomal membrane targeted photosensitizer by fusion of KillerRed to the C-terminus of LAMP1. LAMP1-KillerRed induces lysosomal injury via local production of reactive oxygen species following 561-nm laser irradiation (Bulina et al., 2006). KillerRed activation resulted in the accumulation of eGFP-p62 on lysosomes marked by LAMP2-BFP (Figure 1J,K) (Supplementary Video 1). In regions of cells where KillerRed is inactive, we observe no change in eGFP-p62 intensity, whereas following laser irradiation, eGFP-p62 intensity increased significantly beginning at ten minutes post-laser irradiation (Figure 1L). In addition, we observe limited fluorescence recovery of LAMP1-KillerRed in regions that were irradiated, suggesting the accumulation of eGFP-p62 is specific to damaged lysosomes (Figure 1M). Together these results demonstrate that p62 responds to both cellular-scale lysosomal rupture as well as focal lysosomal damage induced by oxidative stress in HeLa cells.

### p62 responds to lysosomal damage in human and rat neurons

Lysosomal dysfunction has been implicated in several neurodegenerative diseases, and mutations in p62/SQSTM1 are casual for familial ALS (Teyssou et al., 2013). Therefore, we investigated the association of p62 with damaged lysosomes in neurons. We utilized a well-characterized human iPSC line expressing a doxycycline-inducible neurogenin 2 cassette, allowing for efficient differentiation into glutamatergic cortical neurons (i^3^Neurons) (Boecker et al., 2020; Fernandopulle et al., 2018). We treated i^3^Neurons with either ethanol or 1.0mM LLOMe for 2hrs. Using immunocytochemistry, we observed a significant increase in lysosomal area occupied by endogenous p62 in LLOMe-treated i^3^Neurons (EtOH: 4.6 ± 0.49%; LLOMe: 20 ± 1.7%), with no overall change in lysosomal area as compared to control cells (Figure 2A-C).

**Figure 2.**
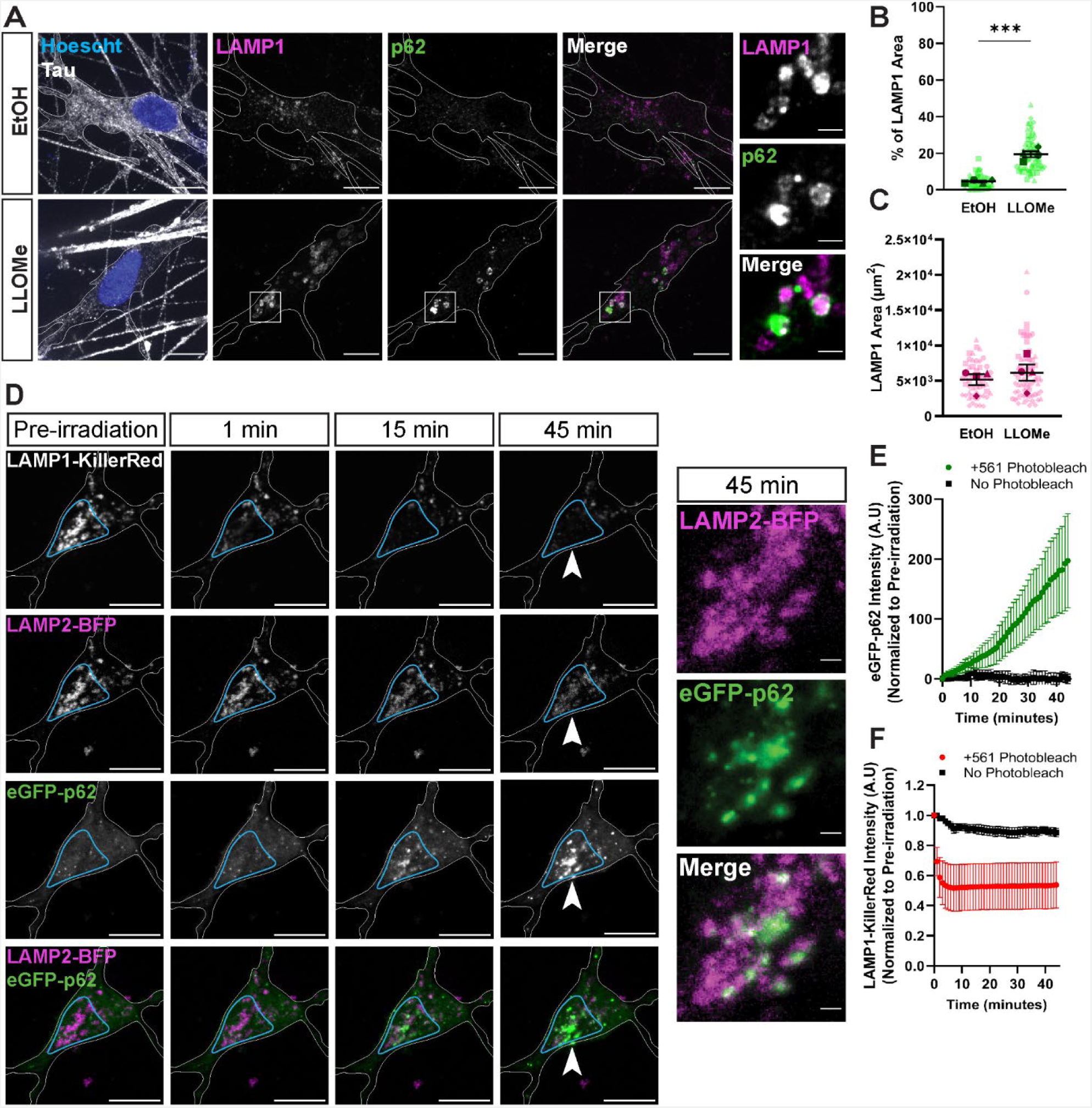
p62 responds to lysosomal damage in human and rat neurons. (A) Representative max projections of anti-p62 in human iPSC-derived neurons (i^3^Neurons), which were treated with EtOH or 1.0mM LLOMe for 2 hours. Cell outlines are traced. White boxes indicate inset image region. Inset images are single z-plane images, demonstrating p62 ring-like structures. Scale Bar: 10µm; inset: 2µm. (B) Quantification of the overlapping area from max projections between p62 and LAMP1 in the different conditions (Unpaired t-test, p < 0.001, N=4, n_(EtOH)_= 54, n_(LLOMe)_=65). (C) Quantification of LAMP1 area from max projections in the different conditions (Unpaired t-test, p=0.5047, N=4, n_(EtOH)_= 54, n_(LLOMe)_=65). (D) Representative images of rat hippocampal neurons transiently transfected with LAMP1-KillerRed, LAMP2-BFP, and eGFP-p62. Images depict pre- and post-561nm laser irradiation. Cell outlines are traced. Blue boxes reflect regions irradiated. Arrowheads reflect location of inset images at 45min post-laser activation. Scale Bar: 10µm; inset: 2µm. (E) Quantification of eGFP-p62 intensities over 45 minutes within regions either activated by laser irradiation or untreated. (F) Quantification of LAMP1-KillerRed intensities over 45 minutes within regions either activated by laser irradiation or untreated.

p62 surrounded lysosomes, appearing as rings in single plane images, indicating a potential role for p62 in the clearance of damaged lysosomes (Figure 2A). We assayed the recruitment of other selective autophagy receptors, including NDP52, Optineurin, and TAX1BP1. We observed a small but significant enrichment of TAX1BP1 on lysosomes following LLOMe treatment in accordance with a previous report (EtOH: 2.7 ± 0.30%; LLOMe: 8.5 ± 1.8%) (Supplemental Figure 1A,B) (Eapen et al., 2021). However, we observed no enrichment of NDP52 (EtOH: 3.5 ± 1.7%; LLOMe: 5.7 ± 0.54%) or Optineurin (EtOH: 3.9 ± 0.29%; LLOMe: 4.9 ± 0.71%) (Supplemental Figure 1C-F).

The comparison of the recruitment of endogenous receptors suggests a key role for p62 in lysosomal quality control in neurons, so we next investigated the temporal and spatial specificity of p62 recruitment. We transiently expressed LAMP1-KillerRed for 48hrs in rat hippocampal neurons. KillerRed activation induced eGFP-p62 accumulation on lysosomes labeled by LAMP2-BFP (Figure 2D-F). Moreover, eGFP-p62 accumulation only occurred in regions that received 561nm-laser irradiation, beginning five minutes post-irradiation (Figure 2E) (Supplementary Video 2). Thus, p62 also responds rapidly to focally-induced lysosomal damage in neurons.

### LAMTOR2 is a lysosomal damage reporter in HeLa cells and neurons

Many studies to date have focused on the recruitment of galectins to monitor lysosomal damage (Aits et al., 2015; Eapen et al., 2021; Jia et al., 2020a; Liu et al., 2020a; Maejima et al., 2013; Papadopoulos et al., 2017; Radulovic et al., 2018; Skowyra et al., 2018). Galectins are cytosolic proteins capable of interacting with B-galactose-containing carbohydrates, and these carbohydrates within the lysosomal glycocalyx are exposed following rupture, driving galectin recruitment to the damaged lysosome (Aits et al., 2015; Barondes et al., 1994; Maejima et al., 2013; Paz et al., 2010). Galectin-1, -3, -8, and -9 have been demonstrated to localize to damaged lysosomes, although galectin-3 is the most commonly used reporter of lysosomal damage because of its rapid response time in human cell lines (Aits et al., 2015; Radulovic et al., 2018; Skowyra et al., 2018)

Transcriptomic analysis predicts that these four galectins are expressed at low levels in the human central nervous system (Aits et al., 2015). We used immunoblotting to evaluate the expression of galectin-3 in i^3^Neurons and mouse cortical neurons, as compared to HeLa cells. In contrast to the robust expression of galectin-3 in HeLa cells, we observed no detectable expression of galectin-3 on western blots of cell lysates from either mouse cortical neurons or human i^3^Neurons (Figure 3A-B). In comparison, galectin-8 was expressed in both mouse and human neurons, and may therefore be a more relevant reporter of neuronal lysosomal damage in these cell types (Figure 3A,C). However, galectin-8 responds to endolysosomal rupture following *Salmonella enterica* serovar Typhimurium (*S*. Typhimurium) infection and promotes autophagy via recruitment of NDP52 to damaged vesicles (Thurston et al., 2012). This observation suggests that transient overexpression of galectin-8 may promote lysophagy, potentially obscuring a requirement for p62. Thus, we sought a galectin-independent method to rapidly identify damaged lysosomes.

**Figure 3.**
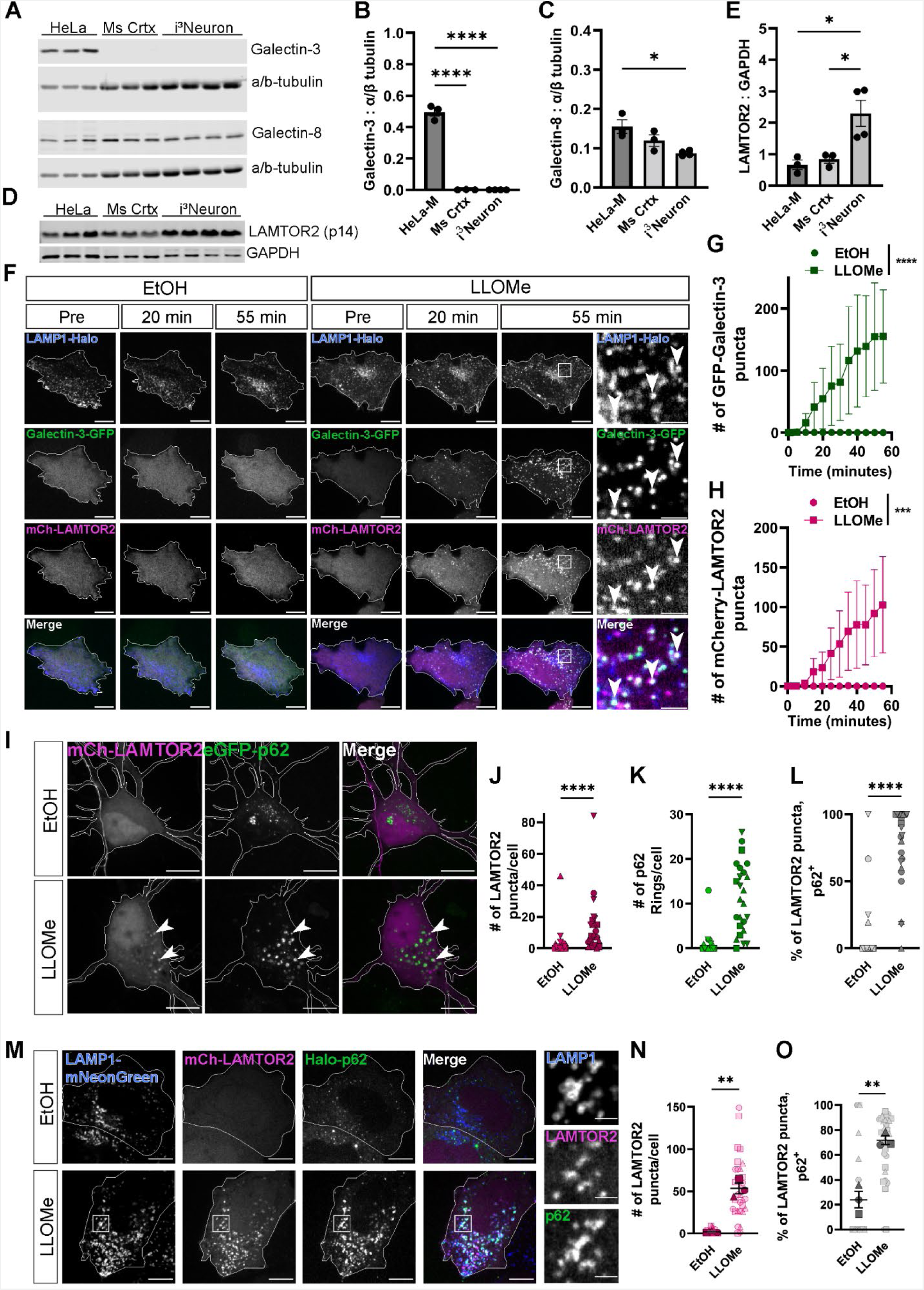
LAMTOR2 is a putative lysosomal damage reporter. (A) Western blot analysis of cell lysates from HeLa cells, mouse cortical neurons (Ms Ctx), or i^3^Neurons. Lysates were evaluated for expression of galectin-3, galectin-8, and α/β-tubulin. (B) Quantification of anti-galectin-3 band intensities following normalization to anti-α/β-tubulin intensity (One-way ANOVA, p_(HeLa/Ms Ctx)_ < 0.0001, p_(HeLa/i3Neuron)_ < 0.001, p_(Ms Ctx/i3Neuron)_ = 0.9988, N=3). (C) Quantification of anti-galectin-8 band intensities following normalization to anti-α/β-tubulin intensity (One-way ANOVA, p_(HeLa/Ms Ctx)_ = 0.1723, p_(HeLa/i3Neuron)_ < 0.05, p_(Ms Ctx/i3Neuron)_ = 0.1883, N=3). (D) Western blot analysis of cell lysates from HeLa cells, Ms Ctx, or i^3^Neurons. Lysates were evaluated for expression of LAMTOR2 (p14) and GAPDH. (E) Quantification of anti-LAMTOR2 band intensities following normalization to anti-GAPDH intensity (One-way ANOVA, p_(HeLa/Ms Ctx)_ = 0.9430, p_(HeLa/i3Neuron)_ < 0.05, p_(Ms Ctx/i3Neuron)_ < 0.05, N=3). (F) Representative images of time course experiment in HeLa cells that were transfected with LAMP1-HaloTag, GFP-Galectin-3, and mCherry-LAMTOR2. Cells were treated with either EtOH or 750µM LLOMe after imaging a pre-treatment frame. Cells were images for 55 minutes following the addition of the drug or vehicle control. Cell outlines are traced. White boxes indicate inset image region. Arrowheads indicate regions of co-localization between LAMP1, Galectin-3, and LAMTOR2. Scale Bar: 10µm; inset: 2µm. (G) Quantification of total GFP-Galectin-3 puncta in cells treated with either EtOH or 750µM LLOMe (Two-way ANOVA, p < 0.0001, N=3). (H) Quantification of total mCherry-LAMTOR2 puncta in cells treated with either EtOH or 750µM LLOMe (Two-way ANOVA, p < 0.001, N=3). (I) Representative max projections of embryonic rat hippocampal neurons (DIV11) that were transiently transfected with mCherry-LAMTOR2 and eGFP-p62. Cells were treated with either EtOH or 1.0mM LLOMe for 2 hours. Cell outlines are traced. Arrowheads indicate examples of mCherry-LAMTOR2 and eGFP-p62 co-localization. Scale Bar: 10µm; inset: 2µm. (J) Quantification of total mCherry-LAMTOR2 puncta in rat hippocampal neurons cells treated with either EtOH or 1.0M LLOMe (Mann-Whitney U test, p < 0.0001, N_EtOH_ = 26, N_LLOMe_ = 25). (K) Quantification of total eGFP-p62 puncta in rat hippocampal neurons cells treated with either EtOH or 1.0M LLOMe (Mann-Whitney U test, p < 0.0001, N_EtOH_ = 26, N_LLOMe_ = 25). (L) Quantification of the percentage of total mCherry-LAMTOR2 puncta co-localizing with eGFP-p62 in rat hippocampal neurons cells treated with either EtOH or 1.0M LLOMe (Mann-Whitney U test, p < 0.0001, N_EtOH_ = 26, N_LLOMe_ = 25). (M) Representative single z-plane images of LAMP1-mNeon HeLa cells. Cells were transiently transfected with mCherry-LAMTOR2 and Halo-p62. Cells were treated with either EtOH or 750µM LLOMe. Cells were live-cell imaged for one hour following the addition of drug or solvent control. Cell outlines are traced. White boxes indicate inset image region. Scale Bar: 10µm; inset: 2µm. (N) Quantification of total mCherry-LAMTOR2 puncta per HeLa cell during treatment protocol (Unpaired t-test, p < 0.01, N=3, n_EtOH_ = 28, n_LLOMe_ = 32). (O) Quantification of the percentage of total mCherry-LAMTOR2 puncta co-localizing with eGFP-p62 per HeLa cell during treatment protocol (Unpaired t-test, p < 0.01, N=3, n_EtOH_ = 28, n_LLOMe_ = 32).

Recent work has demonstrated that LAMTOR2, also referred to as p14, responds to endolysosomal damage following *S.* Typhimurium infection (Lin et al., 2019). LAMTOR2 is a cytosolic component of the Ragulator complex and regulates cellular metabolism (Sancak et al., 2010). Importantly, LAMTOR2 was not required for autophagosome formation following *S.* Typhimurium infection. Overexpression of LAMTOR2 does not disrupt nutrient sensing (Bar-Peled et al., 2012; Sancak et al., 2010; Schweitzer et al., 2015). Together, these data suggest LAMTOR2 could be used to label damaged lysosomes without interfering with cellular responses to damaged lysosomes (Lin et al., 2019).

We first tested whether LAMTOR2 was endogenously expressed in HeLa cells, i^3^Neurons, and mouse cortical neurons, and observed LAMTOR2 expression in all cell types tested (Figure 3D,E). We then assayed the recruitment kinetics of GFP-galectin-3 and mCherry-LAMTOR2 to damaged lysosomes, marked by LAMP1-Halo. Both GFP-galectin-3 and mCherry-

LAMTOR2 were recruited to lysosomes upon addition of 750µM LLOMe, co-localizing with LAMP1-Halo in HeLa cells (Figure 3F) (Supplementary Video 3). GFP-galectin-3 localized to damaged lysosomes within ten minutes of LLOMe addition, in accordance with previous reports (Radulovic et al., 2018; Skowyra et al., 2018) (Figure 3F,G). We observed an accumulation of mCherry-LAMTOR2 on damaged lysosomes approximately fifteen minutes after the addition of 750µM LLOMe (Figure 3F,H). There was no significant difference between the number of mCherry-LAMTOR2 puncta per cell as compared to GFP-galectin-3, suggesting that LAMTOR2 is equivalently capable of rapid translocation to lysosomes (Figure 3G,H).

We next wondered whether p62 co-localized with LAMTOR2 across cell types. We next tested whether eGFP-p62 associates to mCherry-LAMTOR2-postive damaged lysosomes in rat hippocampal neurons. LAMTOR2 formed puncta in neurons following the addition of 1mM LLOMe with the majority of LAMTOR2 puncta (80 ± 9.6%) co-localizing with eGFP-p62 (Figure 3I-L). We also observed that lysosomal damage induced ring-like p62 structures following overexpression, mirroring the response of endogenous p62 following LLOMe treatment (Figure 3K). Next, we assayed LAMTOR2 and p62 co-localization in HeLa cells. In a HeLa stable cell line stably expressing LAMP1-mNeonGreen, we transiently co-expressed mCherry-LAMTOR2 and Halo-p62 (Figure 3M); upon addition of 750µM LLOMe, we observed that the majority of LAMTOR2-positive puncta (72 ± 3.5%) co-localized with p62 (Figure 3M-O). Given that LAMTOR2 is actively recruited to damaged lysosomes in both neurons and HeLa cells, we concluded that LAMTOR2 is an effective lysosomal damage reporter to test both the necessity and sufficiency of p62 in lysophagy.

### p62 is necessary and sufficient for lysophagy

p62 belongs to a class of ubiquitin-dependent selective autophagy receptors. These receptors engage core autophagy machinery as well as the specific cargo to be degraded (Farré and Subramani, 2016; Johansen and Lamark, 2020). As noted above, p62 is recruited within 5-10 min to damaged lysosomes (Figure 1J, Figure 2E).

If p62 is critical in lysophagy, we expect p62 to co-localize with autophagy proteins. Importantly, we observed that following lysosomal damage, p62 co-localized with the Atg8-family proteins GABABRAP/L1/L2, referred herein collectively as GABARAP (Supplemental Figure 2A,B). In single z-slices, GABARAP can appear as puncta or as rings where rings are indicative of formed autophagosomes. We observed that 94 ± 0.49% of GABARAP rings were co-positive for p62, further supportive of a role for p62 in lysophagy (Supplemental Figure 2C). In addition, we observed a significant increase in the overlap of lysosomes and p62, with no change in LAMP1 area per cell (Supplemental Figure 2D,E).

Selective autophagy receptors can function both cooperatively and redundantly (Kirkin et al., 2009; Lazarou et al., 2015). To assay for sufficiency, we utilized a well-characterized HeLa cell line, lacking five autophagy receptors (Optineurin, TAX1BP1, NDP52, NBR1, and p62) (referred to as pentaKO HeLa) (Lazarou et al., 2015). We expressed either eGFP-p62, eGFP-TAX1BP1, eGFP-NBR1, or GFP vector control. To assay changes in lysophagy, we evaluated the recruitment of the Atg8-family protein LC3b to damaged lysosomes using immunofluorescence. We assayed LC3b intensity at mCherry-LAMTOR2 puncta following 2hr treatment with 750µM LLOMe. Strikingly, only expression of eGFP-p62 was sufficient to rescue LC3b intensity in pentaKO HeLa cells (p62: 51% increase LC3b intensity; NBR1: 6.5% decrease LC3b intensity; TAX1BP1: 8.9% increase LC3b intensity) (Figure 4A-B, Supplemental Figure 2F). We did not observe changes in LAMTOR2 recruitment across conditions, indicating that lysosomal damage was induced to a similar extent (Supplemental Figure 2G,H) nor did we observe differences in eGFP intensity, consistent with equivalent levels of expression of each of the eGFP-labeled receptors in these assays (Supplemental Figure 2I).

**Figure 4:**
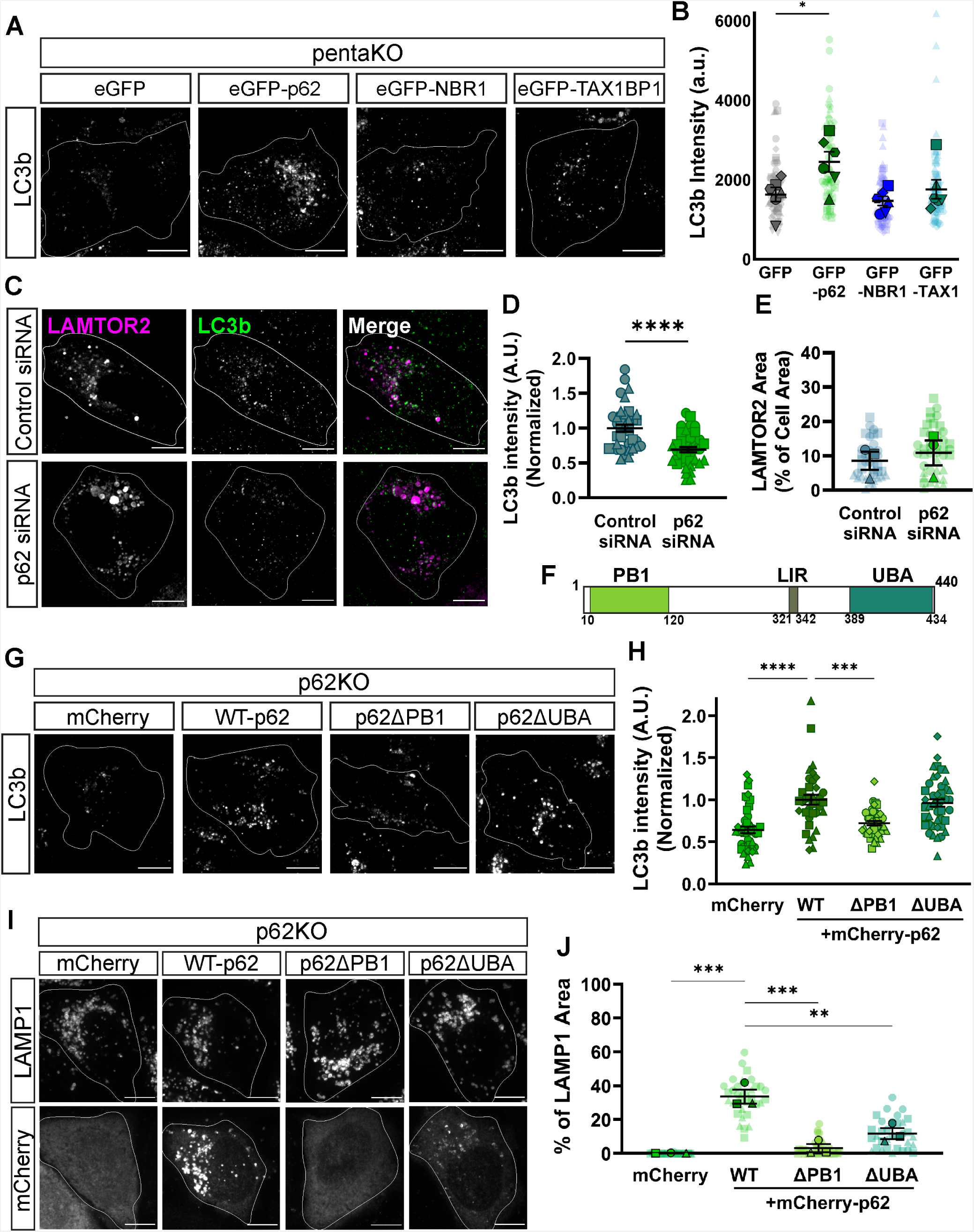
p62 is necessary and sufficient for lysophagy. (A) Representative images of anti-LC3b in pentaKO HeLa cells. Cells were treated with 750µM LLOMe for two hours. Cells were transfected with mCherry-LAMTOR2 and either GFP vector, eGFP-p62, eGFP-NBR1, or eGFP-TAX1BP1. Cell outlines are traced. Scale Bar: 10µm. (B) Quantification of LC3b intensity at LAMTOR2 puncta across different conditions (One-way ANOVA, p_(GFP/p62)_ < 0.05, p_(GFP/NBR1)_ = 0.9042, p_(GFP/TAX1)_ = 0.9417, N=6, n_GFP_ = 72, n_p62_ = 64, n_NBR1_ = 77, n_TAX1_ = 74). (C) Representative images of mCherry-LAMTOR2 and anti-LC3b in HeLa cells, which were transiently transfected with control siRNA or p62 siRNA alongside mCherry-LAMTOR2. HeLa cells were treated with 750µM LLOMe for two hours. Cells were transfected with mCherry-LAMTOR2 and either GFP vector, eGFP-p62, eGFP-NBR1, or eGFP-TAX1BP1. Cell outlines are traced. Scale Bar: 10µm. (D) Quantification of normalized LC3b intensity at LAMTOR2 puncta across different conditions (Mann-Whitney U test, p < 0.0001, N_Control_ = 40, N_p62_ = 41). (E) Quantification of mCherry-LAMTOR2 area across different conditions (Unpaired t-test, p = 0.6338, N=3, n_Control_ = 40, n_p62_ = 41). (F) Schematic of p62 domain architecture, highlighting the N-terminal Phox and Bem1p domain (PB1), LC3-interacting region (LIR), and the ubiquitin-associated domain (UBA). (G) Representative images of anti-LC3b in p62KO HeLa cells, which were transiently transfected with Flag-LAMTOR2 and either mCherry vector, WT-p62, p62ΔPB1, or p62ΔUBA. Cells were treated with 750µM LLOMe for two hours. Cell outlines are traced. Scale Bar: 10µm. (H) Quantification of normalized LC3b intensity at LAMTOR2 puncta across different conditions (Kruskal-Wallis test, p_(WT/mCherry)_ = 0.6631, p_(WT/ΔPB1)_ = 0.1036, p_(WT/ΔUBA)_ = 0.9073, N=4, n_mCherry_ = 44, n_WT_ = 36, n_ΔPB1_ = 39, n_ΔUBA_ = 44). (I) Representative images of anti-LAMP1 and either mCherry vector WT-p62, p62ΔPB1, or p62ΔUBA in p62KO HeLa cells. Cells were treated with 750µM LLOMe for two hours. Cell outlines are traced. Scale Bar: 10µm. (J) Quantification of LAMP1 area overlapping with p62 area across different conditions (One-way ANOVA, p_(WT/mCherry)_ < 0.01, p_(WT/ΔPB1)_ < 0.001, p_(WT/ΔUBA)_ < 0.001, N=3, n_mCherry_ = 33, n_WT_ = 34, n_ΔPB1_ = 34, n_ΔUBA_ = 33).

We next evaluated the specific requirement for p62 in lysophagy. Using transient siRNA knockdown, we achieved 94 ± 0.77% knockdown of p62 (Supplemental Figure 3A,B). We evaluated whether knockdown of p62 prevented lysophagy, measuring the fluorescence intensity of LC3b at LAMTOR2 puncta. Loss of p62 resulted in a significant decrease in LC3b fluorescence intensity (31 ± 3.7% decrease) at LAMTOR2 puncta following 2hrs of 750µM LLOMe treatment (Figure 4C,D), demonstrating a specific requirement for p62 in lysophagy. We confirmed that LAMTOR2 area was unchanged between control and p62 knockdown conditions, suggesting that reduced LC3b recruitment was not due to changes in LAMTOR2 recruitment (Figure 4E). Together, these data demonstrate that p62 is both necessary and sufficient for lysophagy. Moreover, these data suggest that p62 has a unique, non-redundant function in lysophagy that cannot be performed by either TAX1BP1 or NBR1.

To characterize this unique role of p62 in lysophagy, we evaluated which domains of p62 are required (Figure 4F). p62 interacts with ubiquitinated cargo via its C-terminal ubiquitin-associated domain (UBA) (Seibenhener et al., 2004) and with Atg8-family proteins within the developing phagophore via its LC3-interacting region (LIR). The LIR is an evolutionarily conserved motif, consisting of three acidic residues followed by a tryptophan (Johansen and Lamark, 2011). In p62, the LIR consists of DDDW, and mutation of these residues to alanines prevent p62 from interacting with Atg8-family proteins (Pankiv et al., 2007; Turco et al., 2019; Wurzer et al., 2015). p62 also has a N-terminal Phox and Bem1 domain (PB1) that drive p62 homo- or hetero-oligomerization (Lamark et al., 2003; Wilson et al., 2003).

In p62 knockout HeLa cells (p62KO), we transiently expressed FLAG-LAMTOR2 alongside either mCherry-wild type p62 (WT), mCherry-p62ΔPB1 (ΔPB1), mCherry-p62ΔUBA (ΔUBA), or mCherry as a vector control. Immunostaining for LC3b indicated a significant increase in LC3b intensity in cells expressing WT-p62 as compared to p62KO cells expressing the mCherry vector control (36 ± 4.0 % increase) (Figure 4 G,H). Under these conditions, we observed no significant difference in LC3b intensity at LAMTOR2 puncta between WT-and ΔUBA-expressing p62KO cells. We also noted that LC3b intensity was significantly lower in the ΔPB1 condition as compared to cells expressing WT-p62 (28 ± 2.6% decrease) (Figure 4G,H; Supplemental Figure 3C). We observed no differences in LAMTOR2 area, and we observed no differences in mCherry fluorescence among the different p62 constructs (Supplemental Figure 3D,E) These results suggest that the PB1 domain of p62 is crucial for its role in lysophagy, while the UBA domain does not appear to be required under these conditions

We performed analogous experiments in HeLa cells following transient p62 knockdown. We observed similar results in which knockdown of p62 and expression of either mCherry-alone or the ΔPB1 construct both failed to rescue LC3b intensity at LAMTOR2 area, as compared to control cells (mCherry: 28% decrease) (ΔPB1: 25% decrease) (Supplemental Figure 3F,G). We observed no significant differences in FLAG-LAMTOR2 area per cell and no significant differences in mCherry fluorescence intensity among the WT, ΔUBA, and ΔPB1 conditions (Supplemental Figure 3H,I). Together, these data demonstrate that p62 requires its oligomerization domain (PB1) to regulate lysophagy.

p62 is established as an autophagy receptor in the selective clearance of proteins via aggrephagy (ubiquitinated proteins), the selective autophagy of ubiquitinated proteins. In aggrephagy, p62 facilitates autophagosome formation via recruitment of FIP200, a component of the initiation complex. p62-mediated FIP200 recruitment results in the generation of phosphatidylinositol 3-phosphate, which in turn induces the recruitment of WIPI2 (Turco et al., 2019, 2020). Therefore, we assayed whether p62 is required for Halo-WIPI2b puncta formation following 2hrs of 750µM LLOMe. We observed that p62 knockdown resulted in significantly fewer WIPI2b puncta as compared to control siRNA (61% decrease) (Supplemental Figure 4A,B). Rescue of p62 knockdown with WT, ΔUBA, or mCherry-p62-LIR^AAAA^ was able to increase the number of WIPI2b puncta to levels seen with control siRNA. Only the ΔPB1 construct was unable to rescue WIPI2b puncta formation (48% decrease) (Supplemental Figure 4A,B). Again, we observed no changes in cell area or in mCherry fluorescence across conditions (Supplemental Figure 4C,D). These data suggest that lysophagy requires oligomeric p62 early in the autophagy cascade.

We next investigated the mechanism by which p62 oligomerization promotes lysophagy. In p62KO HeLa cells, we evaluated the recruitment of WT, ΔUBA, and ΔPB1 to lysosomes following damage with 750µM LLOMe (Figure 4I). We observed that both ΔUBA and ΔPB1 displayed reduced localization to damaged lysosomes, where WT p62 was recruited to 34 ± 4.1% of lysosomes, ΔUBA recruited to 12 ± 3.2%, and ΔPB1 recruited to 3.0 ± 2.4% (Figure 4I,J). We observed no significant differences in LAMP1 area per cell and no significant differences in mCherry fluorescence intensity between WT, ΔUBA, and ΔPB1 conditions (Supplemental Figure 5A,B). Therefore, loss of oligomerization prevents p62 localization to damaged lysosomes, and this loss of p62 at damaged lysosomes confers a significant deficit in lysophagy. However, lysophagy can proceed with only partial localization of p62, as in the case of the ΔUBA construct.

We were surprised that p62ΔUBA could localize to lysosomes following LLOMe-induced damage, since this construct lacks the well-characterized ubiquitin-binding domain. Thus, we asked whether the localization of ΔUBA was dependent on another selective autophagy receptor. We expressed either WT, ΔUBA, ΔPB1, or a mCherry vector control in pentaKO HeLa. We observed that ΔUBA lost its ability to localize to damaged lysosomes in the pentaKO HeLa (2.0 ± 0.92% of Lysosomes, ΔUBA+), as compared to the partial localization observed in p62KO cells (Supplemental Figure 5C,D). We confirmed that there were no changes in mCherry fluorescence or in LAMTOR2 area across the different p62 constructs, suggesting equivalent lysosomal damage and expression (Supplemental Figure 5E,F). These results suggest that when the UBA domain of p62 is impaired, p62 can cooperatively localize to damaged lysosomes. NBR1 has a similar domain architecture to p62, including a PB1 domain. Thus, we transiently co-expressed ΔUBA with either eGFP-NBR1 or a GFP vector control. We observed an increase in ΔUBA puncta formation, indicating recruitment to damaged lysosomes (Supplemental Figure 5G,H). These results suggest that p62 can still promote lysophagy in the absence of its ubiquitin-binding via interactions with other selective autophagy receptors like NBR1.

In all, these data demonstrate that p62 is a necessary and sufficient receptor for the selective autophagy of damaged lysosomes. p62 requires its oligomerization domain to localize to damaged lysosomes, and loss of p62 localization diminishes lysophagy. However, p62 recruitment is not entirely dependent on direct binding to ubiquitin moieties on damaged lysosomes, as we saw partial localization and rescue of lysophagy that is likely dependent on an association of p62 with the related receptor NBR1. Finally, loss of p62 significantly decreases WIPI2b puncta formation, suggesting, as has been shown in other contexts (Turco et al., 2019), p62 is involved in the recruitment of FIP200 to the damaged lysosome to initiate autophagosome formation in lysophagy.

### HSP27 is recruited to sites of active lysophagy

p62 oligomers are capable of undergoing liquid-liquid phase separation to form p62 condensates (Jakobi et al., 2020; Lamark et al., 2003; Sun et al., 2018; Wilson et al., 2003). p62 condensates can incorporate ubiquitinated cargo as well as autophagy machinery, thus it has been suggested that p62 facilitates selective autophagy of aggregated proteins via condensation (Sun et al., 2018; Turco et al., 2019, 2020). However, it remains an open question whether p62 condensates have a significant role in organellophagy, particularly because p62 is not required for mitophagy (Narendra et al., 2010; Wong and Holzbaur, 2014).

We hypothesized that p62 condensation has a significant role in lysophagy, and proteins that interact with the PB1 domain of p62 may regulate p62 condensates in this process. We searched for proteins that can interact with the PB1 domain of p62 in publicly available lysophagy data sets. Recently, the small heat shock protein HSP27 (also referred to as HSPB1) has been shown in proximity of damaged lysosomes, using APEX2-based proteomics (Koerver et al., 2019). Importantly, HSP27 interacts with p62 via the PB1 domain (Haidar et al., 2019). HSP27 has also been shown to incorporate into FUS and TDP-43 condensates, and is crucial for preventing the fibrilization of FUS and TDP-43 droplets (Liu et al., 2020b; Lu et al., 2021). Therefore, we hypothesized that HSP27 interacts with the PB1 domain of p62 to regulate the ability of p62 to form liquid-like condensates.

HSP27 is phosphorylated at Serine 15, 78, and 82 in response to specific stress conditions, including oxidative stress and hyperosmotic stress (Landry et al., 1992; Niswander and Dokas, 2006). We tested whether lysosomal damage induced by LLOMe was sufficient to induce HSP27 phosphorylation. We treated HeLa cells with EtOH or 750µM LLOMe for 2hrs before evaluating HSP27 phosphorylation via western blot. We observed a significant decrease in total HSP27 but a robust, significant increase in HSP27 phosphorylation following LLOMe treatment (Figure 5A-C), suggesting HSP27 is dynamically responding to lysosomal damage.

**Figure 5:**
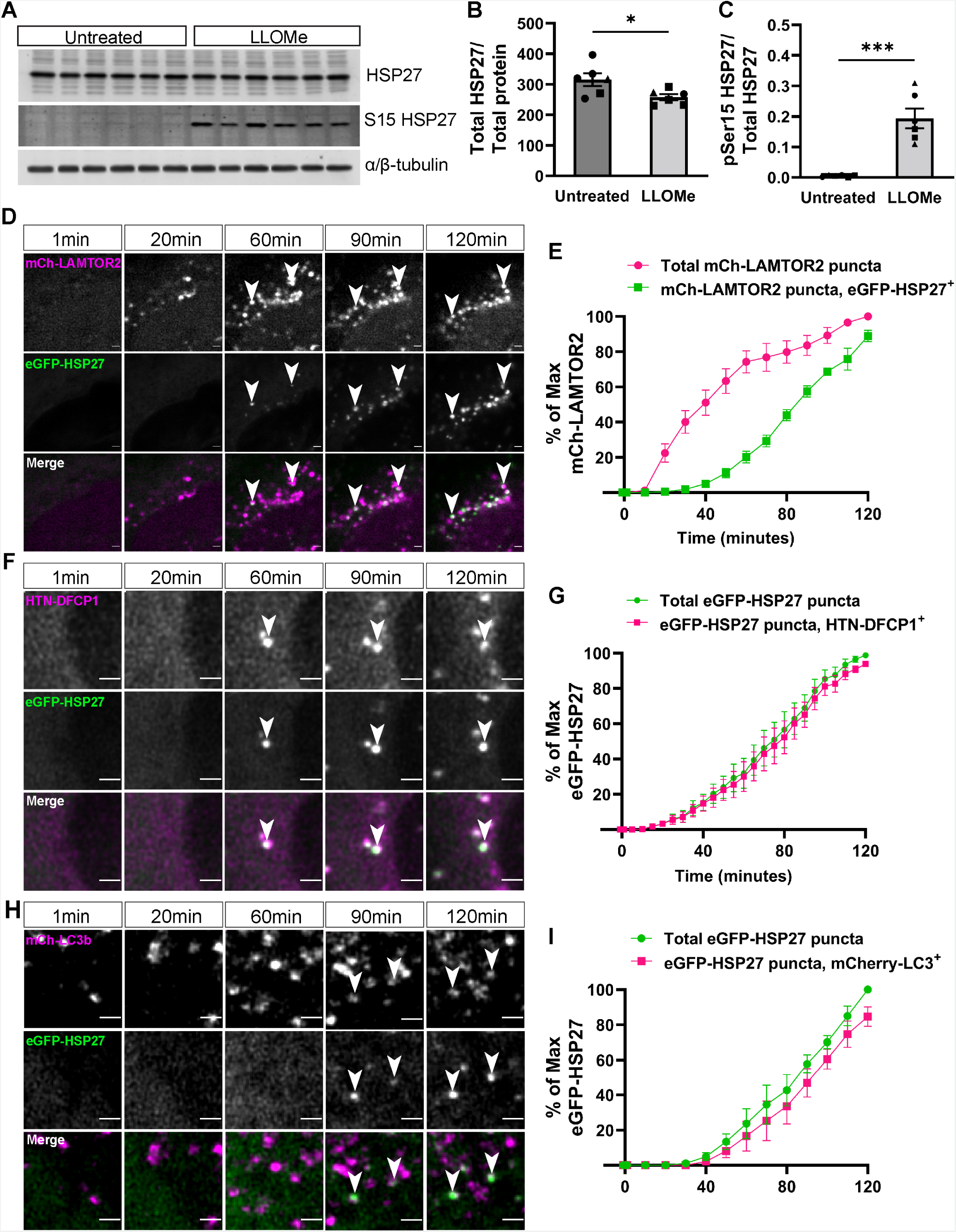
HSP27 is recruited to sites of active lysophagy. (A) Western blot analysis of cell lysates from HeLa cells that were treated with EtOH or 750µM LLOMe for two hours. Lysates were evaluated for expression of HSP27, phospho-S15 HSP27, and α/β-tubulin. (B) Quantification of anti-HSP27 band intensities following normalization to anti-α/β-tubulin intensity (Unpaired t-test, p < 0.05, N=6). (C) Quantification of anti-phospho-S15 HSP27 band intensities following normalization to anti-HSP27 intensity (Unpaired t-test, p < 0.001, N=6). (D) Representative images of mCherry-LAMTOR2 and eGFP-HSP27 puncta in HeLa cells. Cells were treated with 750µM LLOMe after imaging a pre-treatment frame. Cells were imaged for two hours following the addition of LLOMe. Arrowheads indicate co-localization between LAMTOR2 and HSP27. Scale Bar: 2µm. (E) Quantification of total mCherry-LAMTOR2 puncta and the fraction of mCherry-LAMTOR2 puncta that co-localize with eGFP-HSP27 in HeLa cells (Two-way ANOVA, p < 0.001, N=7). (F) Representative images of HaloTag-DFCP1 and eGFP-HSP27 puncta in HeLa cells. Cells were treated with 750µM LLOMe after imaging a pre-treatment frame. Cells were imaged for two hours following the addition of LLOMe. Arrowheads indicate co-localization between DFCP1 and HSP27. Scale Bar: 2µm. (G) Quantification of total eGFP-HSP27 puncta and the fraction of eGFP-HSP27 puncta that co-localize with HaloTag-DFCP1 in HeLa cells (Two-way ANOVA, p = 0.1160, N=7). (H) Representative images of mCherry-LC3b and eGFP-HSP27 puncta in HeLa cells. Cells were treated with 750µM LLOMe after imaging a pre-treatment frame. Cells were imaged for two hours following the addition of LLOMe. Arrowheads indicate co-localization between LC3b and HSP27. Scale Bar: 2µm. (I) Quantification of total eGFP-HSP27 puncta and the fraction of eGFP-HSP27 puncta that co-localize with mCherry-LC3b in HeLa cells (Two-way ANOVA, p < 0.01, N=5).

We next tested whether HSP27 localizes to damaged lysosomes. We transiently co-expressed mCh-LAMTOR2 and eGFP-HSP27 for 24hrs before treating HeLa cells with 750uM LLOMe and simultaneously live-cell imaging. We observed LAMTOR2 forming puncta following ∼20min of lysosomal damage, whereas HSP27 was recruited to LAMTOR2 puncta following ∼40min of lysosomal damage (Figure 5D; Supplementary Figure 6A,B) (Supplementary Video 4). Following two hours of LLOMe, 89 ± 3.2% of LAMTOR2 puncta were co-labeled by eGFP-HSP27, suggesting that HSP27 is recruited to the majority of damaged lysosomes (Figure 5E). Importantly, the timing of HSP27 localization corresponds to previous reports describing the activation of lysophagy (Radulovic et al., 2018).

Next, we wondered whether HSP27 responds similarly to lysosomal damage in i^3^Neurons. We transiently expressed eGFP-HSP27 in i^3^Neurons for 72hrs before treating with 1mM LLOMe for 2hrs. We observed that addition of LLOMe induced HSP27 puncta formation in i^3^Neurons (Supplementary Figure 6C). Together, these data show that HSP27 is capable of responding to lysosomal damage in both HeLa cells and i^3^Neurons.

We next assayed whether HSP27 localized to lysosomes undergoing lysophagy. We first evaluated the co-localization between eGFP-HSP27 and HaloTag-DFCP1. DFCP1, or double FYVE domain-containing protein 1, marks PtdIns(3)P-rich extensions of the endoplasmic reticulum from which autophagosomes originate (Axe et al., 2008). We treated HeLa cells with 750µM LLOMe while live-cell imaging. We observed striking co-localization between HaloTag-DFCP1 and eGFP-HSP27 following ∼50min of lysosomal damage (Figure 5F,G) (Supplementary Video 5). 94 ± 1.4% of HSP27 puncta co-localized with DFCP1 after 2hrs LLOMe (Figure 5G). We then evaluated the co-localization between eGFP-HSP27 and mCherry-LC3b. Using live-cell imaging, we observed that the majority of HSP27 puncta co-localized with mCherry-LC3b. 85 ± 5.6% of HSP27 puncta co-localized with LC3b after 2hrs LLOMe (Figure 5H-I). These results indicate that HSP27 is present at sites of lysophagy, co-localizing with DFCP1 and LC3b. Moreover, as more HSP27 puncta are labeled by DFCP1 compared to the Atg8-family protein LC3b, it is possible that HSP27 recruitment precedes the formation of the phagophore.

### HSP27 regulates p62 condensates and lysophagy

We hypothesized that HSP27 regulates the liquidity of p62 condensates. Thus, we evaluated the association of eGFP-HSP27 with endogenous p62 following lysosomal damage. We transiently expressed eGFP-HSP27 for 24hrs before treating HeLa cells with 750µM LLOMe for 2hrs (Supplemental Figure 6D). We observed significant increases in the overlap of HSP27 with LAMP1 and p62 following lysosomal damage (Supplemental Figure 6E,F). We simultaneously observed increased overlap of p62 with LAMP1, with no differences in LAMP1 area (Supplemental Figure 6G,H). To test the recruitment kinetics of HSP27 and p62, we transiently co-expressed eGFP-HSP27 and mCh-p62 before treating cells with 750µM and live-cell imaging. We observed that 96 ± 1.4% of eGFP-HSP27 was co-localized with mCh-p62 (Figure 6A,B) (Supplementary Video 6). Moreover, we observed eGFP-HSP27 accumulate on p62 puncta following lysosomal damage (Figure 6C). These results show that HSP27 dynamically responds to lysosomal damage and may be regulated by p62 accumulation on damaged lysosomes.

**Figure 6:**
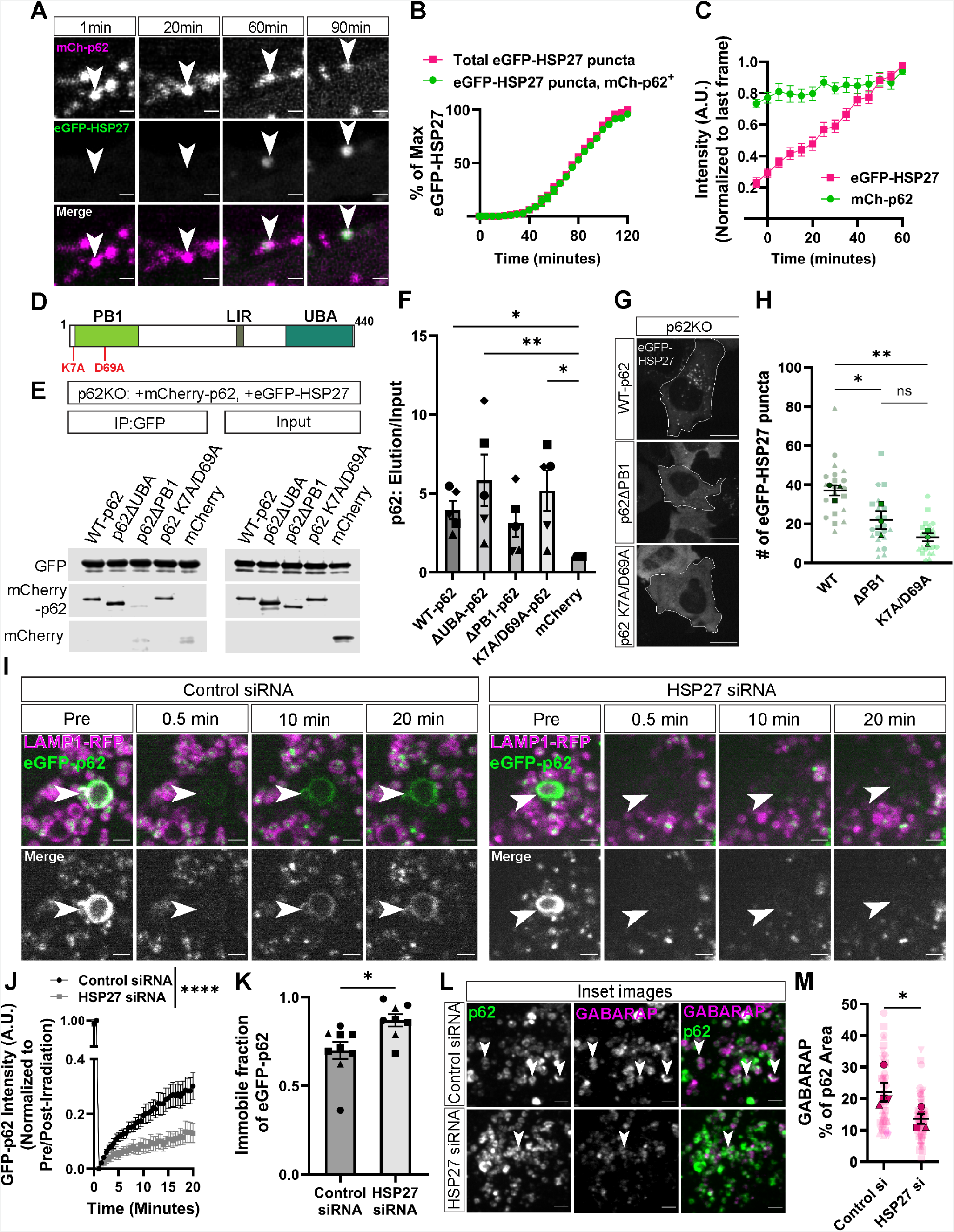
HSP27 regulates p62 condensates. (A) Representative images of mCherry-p62 and eGFP-HSP27 in HeLa cells. Cells were treated with 750µM LLOMe after imaging a pre-treatment frame. Cells were imaged for two hours following the addition of LLOMe. Arrowheads indicate co-localization between p62 and HSP27 over time. Scale Bar: 2µm. (B) Quantification of total eGFP-HSP27 puncta and the fraction of eGFP-HSP27 puncta that co-localize with mCherry-p62 (Two-way ANOVA, p < 0.01, N=4). (C) Quantification of eGFP-HSP27 and mCherry-p62 fluorescence intensities normalized to the fluorescence at last frame analyzed. Puncta were selected if they remained in z-plane for one hour (Two-way ANOVA, p < 0.0001, N=9). (D) Schematic of p62 domain architecture, highlighting K7 and D69, which are key residues in p62 oligomerization. Alanine substitution of these residues prevent homo- and hetero-oligomerization. (E) Representative western blot demonstrating eGFP-HSP27 co-immunoprecipitation of either WT, p62ΔUBA, p62ΔPB1, K7A/D69A p62, or mCherry vector. (F) Quantification of co-immunoprecipitation of mCherry-tagged constructs with eGFP-HSP27 (Kruskal-Wallis test, p_(mCherry/WT-p62)_ < 0.05, p_(mCherry/ΔUBA)_ < 0.01, p_(mCherry/ΔPB1)_ = 0.1922, p_(mCherry/(K7A/D69A))_ < 0.05, N=5). (G) Representative images of eGFP-HSP27 in p62KO HeLa cells that were co-transfected with either WT, p62ΔPB1, or K7A/D69A p62. Cells were treated with 750µM LLOMe for two hours. Cell outlines are traced. Scale Bar: 10µm. (H) Quantification of the number of eGFP-HSP27 puncta per cell (One-way ANOVA, p_(WT/ΔPB1)_ < 0.05, p_(WT/(K7A/D69A))_ < 0.01, p_(ΔPB1/(K7A/D69A)_ = 0.2131, N=3, n_WT_ = 21, n_ΔPB1_ = 22, n _K7A/D69A_ = 23). (I) Representative images of fluorescence recovery of eGFP-HSP27 following photo-bleaching in cells transiently transfected with control siRNA or HSP27 siRNA. Cells were treated with LLOMe for one hour before photo-bleaching, and cells were imaged for 20 minutes following photo-bleaching. Scale Bar: 2µm. (J) Quantification of fluorescence recovery after photo-bleaching of eGFP-p62 (Two-way ANOVA, p < 0.0001, N_Control_ = 9, N_Hspb1_ = 9). (K) Quantification of the immobile fraction of eGFP-p62 across conditions (Unpaired t-test, p < 0.05, N_Control_ = 9, N_Hspb1_ = 8). (L) Representative images of anti-p62 and anti-GABARAP in HeLa cells following transient transfection of control siRNA or HSP27 siRNA. Arrowheads indicate example co-localization events between p62 and GABARAP. Scale Bar: Scale Bar: 2µm. (M) Quantification of the percentage of area of anti-p62 that is occupied by anti-GABARAP (Mann-Whitney test, p < 0.05, N = 4, n_Control_ = 56, n_Hspb1_ = 71).

As HSP27 interacts with the PB1 domain of p62 (Haidar et al., 2019), we next asked whether this interaction depends on the oligomerization of p62. Alanine substitutions of K7 and D69 in the PB1 domain of p62 are sufficient to prevent p62 oligomerization (Figure 6D) (Lamark et al., 2003; Wurzer et al., 2015). Therefore, we immunoprecipitated eGFP-HSP27 and assayed for pull-down of WT-p62, ΔUBA, ΔPB1, K7A/D69A p62, or mCherry-alone in p62KO HeLa cells. We observed increased binding of HSP27 to WT-p62, ΔUBA, and K7A/D69A p62 as compared to the mCherry vector. In contrast, we observed equivalent binding of HSP27 to p62ΔPB1 and mCherry vector (Figure 6E,F; Supplemental Figure 6I). Additionally, HSP27 was immunoprecipitated equivalently across conditions, and we observed equivalent binding to HSP90, a known interactor of HSP27, across conditions (Supplemental Figure 6J,K) (Yang et al., 2015). These results suggest that the interaction of HSP27 and p62 does not directly depend on p62 oligomerization.

Loss of the PB1 domain prevented p62 from localizing to damaged lysosomes (Figure 4I,J). Similarly, both the p62ΔPB1 and the K7A/D69A mutant displayed reduced localization to *S. Typhimurium* in HeLa cells (Wurzer et al., 2015). We wondered whether loss of p62 oligomerization affected HSP27’s response to lysosomal damage. We transiently transfected p62KO HeLa with eGFP-HSP27 and either WT-p62, ΔPB1, or K7A/D69A p62. We treated HeLa cells with 750µM LLOMe for 2hrs before fixation. We then quantified the number of eGFP-HSP27 puncta per cell. We observed significantly fewer HSP27 puncta in p62KO HeLa expressing either ΔPB1 or K7A/D69A p62 as compared to WT-p62 (ΔPB1: 41% decrease; K7A/D69A p62: 64% decrease) (Figure 6G,H; Supplemental Figure 6L). We observed no changes in cell area or in eGFP-HSP27 fluorescence intensity (Supplemental Figure 6M,N). These results demonstrate that although HSP27 can interact with K7A/D69A p62, HSP27 recruitment to damaged lysosomes depends on p62 oligomer formation.

The liquidity of selective autophagy cargo regulates the engulfment of the cargo by autophagosomes. Either increased or decreased liquidity of condensates can impair cargo degradation (Yamasaki et al., 2020). Therefore, we tested whether loss of HSP27 affects p62 condensation. To assay shifts in p62 liquidity, we utilized fluorescence recovery after photo-bleaching, as decreased fluorescence recovery indicates a more gel-like p62 assembly (Carisey et al., 2011). We transiently transfected HeLa cells with either control siRNA or HSP27 siRNA alongside eGFP-p62. We observed 88 ± 2.6% knockdown efficiency of HSP27 in HeLa cells (Supplemental Figure 6O,P). We damaged lysosomes with 750µM LLOMe for one hour before photo bleaching eGFP-p62 ring structures and measuring fluorescence recovery (Figure 6I). Depletion of HSP27 resulted in slower fluorescence recovery (13 ± 3.5% recovery at t=20min) as compared to control cells (30 ± 4.8% recovery at t=20min) (Figure 6I,J) (Supplementary Video 7). Similarly, we quantified the immobile fraction of p62 using the fluorescence recovery assay. The immobile fraction reflects the solid-state p62 or fraction of p62 that did not exchange with the cytosolic p62 (Carisey et al., 2011). We also observed a significant increase in the immobile fraction of eGFP-p62 following knockdown of HSP27 (Figure 6K). These results suggest that HSP27 functions to maintain the liquidity of p62 condensates at damaged lysosomes.

We then tested whether loss of HSP27 affects the engulfment of damaged lysosomes marked by p62. We transiently transfected either control siRNA or HSP27 siRNA for 48hrs before treating cells with 750µM LLOMe for two hours. We performed immunocytochemistry to evaluate the association of the Atg8-family protein GABARAP to p62-positive damaged lysosomes. Following LLOMe treatment, we observed a decrease in GABARAP overlapping with p62 in cells depleted of HSP27 (14 ± 1.6%) as compared to control cells (22 ± 2.9%) (36% decrease) (Figure 6L,M; Supplemental Figure 6Q). We also observed a significant decrease in GABARAP fluorescence intensity at p62 segmented area (15 ± 1.3% decrease) (Supplemental Figure 6R). However, we observed no significant decrease in p62 association with lysosomes or in LAMP1 area between conditions (Supplemental Figure 6S,T). These results together demonstrate that HSP27 is crucial for p62-mediated lysophagy and regulates the liquidity of p62 condensates, affecting the ability to locally assemble an autophagosome.

### ALS-associated mutations in p62 perturb lysophagy

Amytrophic Lateral Sclerosis (ALS) is a fatal neurodegenerative disease, targeting the motor neurons of the brain and spinal cord. The majority of ALS cases are sporadic, yet 10% of cases are familial (Milicevic et al., 2022). In ALS patients, motor neurons accumulate proteinaceous inclusions that result in degeneration. p62 is commonly found in these inclusions, and further, p62 has been suggested to promote the formation of these inclusions (Al-Sarraj et al., 2011; Foster et al., 2021; Mitsui et al., 2018). Mutations in p62 have been identified in patients with sporadic and familial ALS (Teyssou et al., 2013). We wondered whether ALS-associated mutations in p62 affect lysophagy, focusing on the well-characterized mutations L341V, P392L, and G425R (Figure 7A). L341V maps to the LIR motif, residing in the mAtg8 binding pocket (Chen et al., 2014; Ichimura et al., 2008). Importantly, the L341V mutation significantly reduces binding of p62 to mAtg8 proteins (Goode et al., 2016). P392 is in the UBA domain of p62 but is remote to the ubiquitin binding pocket. The P392L mutation mildly alters the secondary structure of the UBA domain, yet P392L results in decreased binding of p62 to polyubiquitin and decreased clearance of ubiquitin aggregates (Cavey et al., 2006; Deng et al., 2020; Garner et al., 2011). G425 is within the ubiquitin binding pocket of the UBA domain, and G425R results in complete loss of ubiquitin binding as well as decrease in aggregate clearance (Deng et al., 2020; Garner et al., 2011).

**Figure 7:**
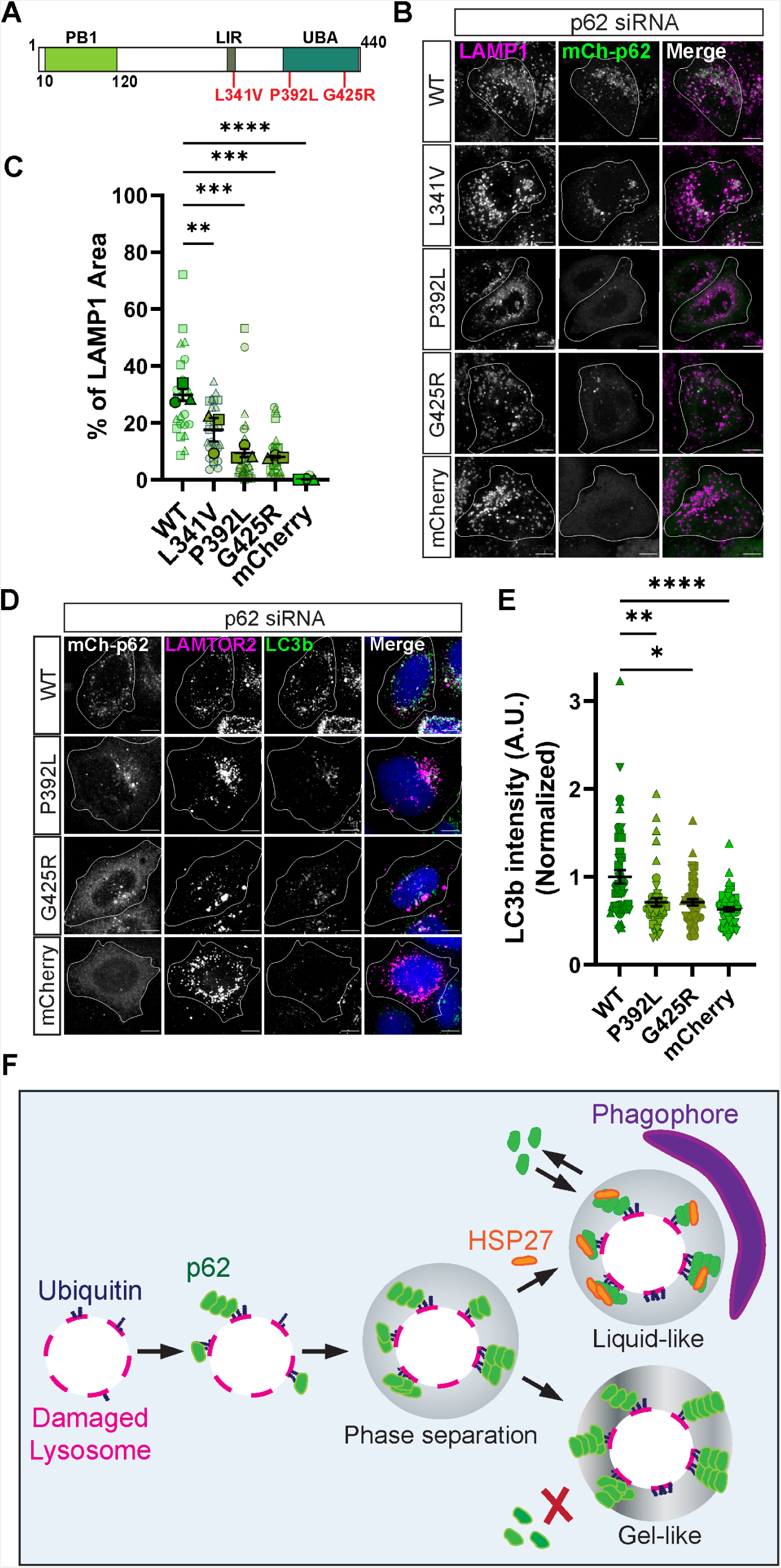
ALS-associated mutations in p62 perturb lysophagy. (A) Schematic of p62 domain architecture, highlighting specific ALS-associated mutations (L341V, P392L, G425R). (B) Representative images of anti-LAMP1 in HeLa-M cells that were transiently transfected with p62 siRNA alongside either WT-p62, L341V, P392L, G425R, or mCherry Vector. HeLa-M cells were treated with 750µM LLOMe for two hours. Cell outlines were traced. Scale Bar: 10µm. (C) Quantification of LAMP1 area overlapping with p62 area across different conditions (One-way ANOVA, p_(WT/L341V)_ < 0.01, p_(WT/P392L)_ < 0.001, p_(WT/G425R)_ < 0.001, p_(WT/mCherry)_ < 0.0001, N=3, n_WT_ = 22, n_L341V_ = 27, n_P392L_ = 27, n_G425R_ = 27, n_mCherry_ =25). (D) Representative images of anti-LC3b in HeLa-M cells that were transiently transfected with p62 siRNA alongside Flag-LAMTOR2 and either WT-p62, P392L, G425R, or mCherry Vector. HeLa-M cells were treated with 750µM LLOMe for two hours. Cell outlines were traced. Scale Bar: 10µm. (E) Quantification of normalized LC3b intensity at LAMTOR2 puncta across different conditions (Kruskal-Wallis test, p_(WT/P392L)_ < 0.01, p_(WT/G425R)_ < 0.05, p_(WT/mCherry)_ < 0.0001, N_WT_ = 49, N_P392L_ = 51, N_G425R_ = 51, N_mCherry_ = 49). (F) A model for p62-dependent lysophagy. p62 is recruited to damaged lysosomes through ubiquitin. Accumulation of p62 on damaged lysosomes results in phase separation. This phase separation of p62 is critically regulated by the small heat shock protein HSP27. HSP27 maintains the liquidity of p62 condensates, facilitating engulfment by autophagosomes.

We first assayed the localization of L341V, P392L, or G425R following lysosomal damage. In HeLa cells depleted of p62, we rescued with either WT, L341V, P392L, or G425R p62, observing that L341V (18 ± 4.2%), P392L (9.4 ± 1.4%), and G425R (8.1 ± 0.33%) mutants resulted in reduced localization to LAMP1 vesicles as compared to WT-p62 (29 ± 2.1%) (Figure 7B,C). We observed no changes in LAMP1 area or in mCherry-p62 fluorescence intensity across the conditions (Supplemental Figure 7A,B).

Finally, we assayed whether the P392L or G425R mutants affected mAtg8 levels at damaged lysosomes. These mutants had the strongest mis-localization effect, thus we hypothesized that rescue with either P392L or G425R would decrease lysophagy. Consistent with this hypothesis, in HeLa cells depleted of p62, we observed that P392L (29 ± 4.5% decrease) or G425R (29 ± 3.6% decrease) mutants had reduced LC3b intensity as compared to WT-p62 (Figure 7D,E, Supplemental Figure 7C). Again, we observed no differences in LAMTOR2 area or mCherry fluorescence intensity across the conditions (Supplemental Figure 7D,E). Therefore, ALS-associated mutations that disrupt the ability of p62 to bind ubiquitin lead to reduced lysophagy.

## Discussion

Lysophagy is the primary mechanism by which ruptured lysosomes are degraded. In cell lines, damaged lysosomes have been shown to recruit several autophagy receptors, including NDP52, TAX1BP1, and p62 (Eapen et al., 2021; Koerver et al., 2019). However, the roles of each of these selective autophagy receptors in lysophagy has not been investigated in detail. Here, we define the role of p62 as a selective autophagy receptor in lysophagy (Figure 7F). We identify the robust recruitment of p62 to damaged lysosomes using different lysosomal injury paradigms in both HeLa cells and in neurons. Further, we observe that p62 is both necessary and sufficient for efficient lysophagy.

Strikingly, we found that expression of p62 was sufficient to rescue LC3b intensity at damaged lysosomes in pentaKO HeLa, while expression of either TAX1BP1 or NBR1 was insufficient, suggesting that p62 has a unique, non-redundant function in lysophagy. In a sequential model for the recruitment of autophagy receptors, p62 establishes an autophagy platform that precedes and is required for TAX1BP1-mediated recruitment of the autophagosome initiation complex (Turco et al., 2021). Recently, TAX1BP1 was identified as critical in lysophagy, recruiting the autophagosome initiation complex and the ULK1-activating kinase TBK1 (Eapen et al., 2021). Therefore, in lysophagy, p62 may function upstream of other lysophagy receptors. In support of this model, we observed cross-talk between selective autophagy receptors in lysophagy. Even when the ubiquitin-binding domain is deleted, p62ΔUBA can be co-recruited to damaged lysosomes by overexpression of the selective autophagy receptor NBR1. Thus, selective autophagy receptors can function cooperatively but not redundantly in lysophagy.

We observed that the PB1 domain of p62 plays a critical role in lysophagy. The PB1 domain of p62 facilitates the self-oligomerization of p62, allowing for p62 condensates to form. The significance of p62 condensates in organellophagy has been unclear as they are not required for mitophagy (Danieli and Martens, 2018), but here, we observed encapsulation of lysosomes via p62 and the requirement for the PB1 domain of p62. Thus, we wondered whether p62 condensates are critical in lysophagy. We sought to identify proteins capable of regulating p62 condensates and investigate their role in lysophagy. We identified the small heat shock protein HSP27 dynamically responds to lysosomal damage. More specifically, we observed accumulation of HSP27 on p62 puncta following lysosomal damage, demonstrating that HSP27 may be regulating p62.

HSP27 has been shown to incorporate into FUS and TDP-43 liquid-phase separated condensates (Liu et al., 2020b; Lu et al., 2021). Thus, we hypothesized that HSP27 interacts with p62 to regulate its phase separation. We observe that depletion of HSP27 decreased the liquid phase properties of p62 on damaged lysosomes. Moreover, we observed striking phosphorylation of HSP27 following lysosomal damage. Phosphorylation of HSP27 promotes the incorporation of HSP27 into FUS condensates, preventing FUS aggregation (Liu et al., 2020b). Therefore, following lysosomal damage, HSP27 may be phosphorylated to prevent the aggregation of p62 that accumulates on damaged lysosomes (Figure 7F).

We hypothesized that decreasing the liquidity of p62 condensates would decrease lysophagy. Accordingly, depletion of HSP27 resulted in decreased GABARAP association with p62 at damaged lysosomes. Therefore, HSP27 interacts with p62 on damaged lysosomes, promoting the liquidity of p62 condensates and facilitating lysophagy. The interaction between HSP27 and p62 occurs within the PB1 domain of p62. Of note, this interaction does not require oligomerization of p62, as HSP27 can interact with K7A/D69A p62. Interestingly, HSP27 interacts with FUS via the N-terminal low complexity domain in FUS (Liu et al., 2020b). Within the PB1 domain, p62 has a predicted low complexity region (PlaToLoCo, Accession Number: <u>BC019111.1)</u>, thus it is possible that HSP27 interacts with p62 via weak interactions in this predicted low complexity region.

We also observe robust spatial and temporal co-localization between HSP27 and the autophagy protein DFCP1. These data suggest either that p62 remodeling by HSP27 is fast, eliciting the rapid formation of an autophagosome or that HSP27 recruits an additional factor that elicits autophagosome formation. Further mechanistic work is needed to test this hypothesis.

Interestingly, both HSP27 and p62 have been implicated in neurodegenerative disease. Mutations in HSP27 are causative for distal hereditary motor neuropathy and Charcot-Marie Tooth Disease Type 2F (Evgrafov et al., 2004). HSP27 is also decreased in motor neurons from ALS patients (Lu et al., 2021). Mutations in p62 have been associated with both sporadic and familial cases of ALS (Milicevic et al., 2022). We observe that disease-associated mutations in p62 (L341V, P392L, G425R) decease p62 localization to damaged lysosomes. Consequently, we observe that the P392L and G425R mutants of p62 are unable to rescue LC3b intensity at damaged lysosomes, indicating decreased lysophagy. Several ALS-associated mutations in p62 cause decreased liquidity in p62 condensates (Faruk et al., 2021). Thus, deficits in p62 localization or decreased liquidity in p62 condensates may be involved in pathophysiology of ALS. Importantly, multisystem proteinopathy, which includes the neurodegenerative aspects of ALS, has been linked previously to lysophagy, as disease-causing mutations in Valosin-containing protein result in defective lysophagy (Papadopoulos et al., 2017). Therefore, impaired lysophagy could represent a common characteristic in proteinopathies like ALS.

There have been previous conflicting reports regarding the requirement of p62 in lysophagy. Depletion of p62 resulted in decreased clearance of Galectin-3 puncta, yet deletion of p62 resulted in no change in Galectin-3 acidification, suggesting that deletion of p62 had no effect on lysophagy (Eapen et al., 2021; Papadopoulos et al., 2017). However, here we demonstrate that p62 functions as an essential lysophagy receptor. We suggest that these apparently conflicting reports may result from the methods used to monitor damaged lysosomes. Damaged lysosomes are frequently identified by Galectin-3 recruitment, but in cells overexpressing Galectin-3, p62 was found not to be required for lysophagy (Eapen et al., 2021). Together, these observations suggest that Galectin-3 overexpression may mask the requirement of p62, potentially by enhancing other pathways to autophagosome formation. Consistent with this possibility, galectin-3 is capable of undergoing liquid-liquid phase separation in vitro as well as on endomembranes following LLOMe treatment (Zhao et al., 2021). Therefore, we propose that Galectin-3 overexpression may compensate for loss of p62 condensates. However, the significance of endogenous Galectin-3 phase separation in lysophagy remains to be tested.

In summary, our results demonstrate that p62 functions as a lysophagy receptor. We demonstrate that p62 is both necessary and sufficient for lysophagy. Our results demonstrate that p62 likely serves as a platform for autophagosome biogenesis in lysophagy. This platform requires precise tuning via HSP27, which maintains the liquidity of p62 condensates and prevents the formation of a p62 gel-like condensate (Figure 7F). The interaction between p62 and HSP27 may be conserved in other cellular contexts, as both proteins are known to respond to cellular stress conditions (Haidar et al., 2019; Kumar et al., 2022; Landry et al., 1992; Niswander and Dokas, 2006). Moreover, both p62 and HSP27 previously have been implicated in ALS (Lu et al., 2021; Teyssou et al., 2013). The recent demonstration that another p62-PB1 binding protein, Nur77, regulates the liquidity of p62 condensates in Celastrol-induced mitophagy (Peng et al., 2021), suggests that regulation of p62 condensates via protein-protein interactions in the PB1 domain may reflect a common mechanism across many types of selective autophagy. Further investigation into the relationship between p62 and HSP27 may uncover mechanisms that can be therapeutically targeted to treat proteinopathies like ALS as well as other disorders of lysosomal dysfunction.

## Materials and Methods

### Antibodies

The following primary antibodies were used: rabbit polyclonal anti-LAMP1 (Abcam, ab24170, Lot: GR3409154, WB: 1:1000, IF: 1:500); sheep polyclonal anti-LAMP1/CD107a (R&D) Systems, AF4800, Lot: CAYN022012A, CAYN022107B, CAYN0221081, IF: 1:40); sheep polyclonal anti-LAMP1/CD107a Alex Flour 488-conjugated (IC7985G, Lot: ADNG0420101, ADNG0420041, IF: 1:40); mouse monoclonal anti-p62 (Abcam, ab56146, Lot: GR3367558, GR3397093, IF: 1:500); guinea pig polyclonal anti-p62 (American Research Products, 03-GP62-C, Lot: 703241-03, WB: 1:3000); rabbit polyclonal anti-FIP200 (Proteintech, 17250-1-AP, Lot: 0050110, WB: 1:1000); rabbit monoclonal anti-TAX1BP1 (Abcam, ab176572, Lot: GR137934, IF: 1:200, WB: 1:1000), rabbit monoclonal anti-GABARAP (E1J4E) (Cell Signaling, 13733, Lot: 3, WB: 1:1000); rabbit monoclonal anti-GABARAP+GABARAPL1+GABARAPL2 (Abcam, ab109364, Lot: GR3232141, IF: 1:100); rabbit polyclonal anti-LC3b (Novus Biologicals, NB100-2220, Lot: DP-1, WB: 1:1000); rabbit polyclonal anti-LC3b (Abcam, ab48394, Lot: GR3401900, GR3374546, GR3339674, IF: 1:200); rabbit polyclonal anti-NDP52 (Abcam, ab68588, Lot: GR197350; IF: 1:200); rabbit polyclonal anti-Optineurin (Abcam, ab23666, Lot: GR271296, IF: 1:200); rabbit monoclonal anti-Tau (Abcam, ab8763, Lot: 158781, IF: 1:200); mouse monoclonal anti-Tau (Sigma, MAB3420, Lot: 2876166, IF: 1:1000); mouse monoclonal anti-Galectin-3 (Santa Cruz Biotech, sc-32790, Lot: G1620, IF: 1:1000); rabbit polyclonal anti-Galectin-8 (Abcam, ab42879, Lot: abGR3277211, WB: 1:1000); mouse monoclonal anti-Hsp27 (Santa Cruz Biotech, sc-13132, Lot: I0821, WB: 1:200); rabbit monoclonal anti-pSer15 Hsp27 (Sigma, SAB4300134, Lot: 871511164, WB: 1:500); rabbit monoclonal anti-HSP90 (C45G5) (Cell Signaling, 4877T, Lot: 5, WB: 1:1000); chicken polyclonal anti-mCherry (Abcam, ab205402, Lot: GR3176028, GR3368071, GR3424850, WB: 1:1000, IF: 1:200); chicken polyclonal anti-GFP (Aves Lab, GFP-1020, Lot: GFP3717982, WB: 1:1000; IF: 1:1000); mouse monoclonal anti-GFP (Abcam, ab1218, Lot: GR3392723, IF: 1:200, IP: 1.25µg/condition); mouse monoclonal anti-HA-Tag (16B12) Alexa flour-594 (Invitrogen, A21288, Lot: 2218838, IF: 1:100); mouse monoclonal anti-FLAG M2 (Sigma, Lot: SLBG5673V, SLCF4933,SLCJ4124, SLCK5688, IF: 1:200).

The following secondary antibodies were used for immunofluorescence: AlexaFluor 488 goat anti-rabbit IgG (H+L) (Invitrogen, A11034, Lot: 2256692); AlexaFluor 594 goat anti-rabbit IgG (H+L) (Invitrogen, A11037, Lot: 2307302, 1915919); AlexaFluor 647 goat anti-rabbit IgG (H+L) (Invitrogen, A31573, Lot: 2359136); AlexaFluor 488 goat anti-chicken IgG (H+L) (Invitrogen, A11039, 2079383); AlexaFluor 555 goat anti-chicken IgG (H+L) (Invitrogen, A21437, Lot: 1719681); AlexaFluor 594 goat anti-mouse IgG (H+L) (Invitrogen, A11032, Lot: 2129448); goat anti-mouse IgG (H+L) Alexa Fluor Plus 647 (Invitrogen, A32728, Lot: WK331591, 1826679); AlexaFluor 594 donkey anti-sheep IgG (H+L) (Invitrogen, A11016, Lot: 536045); GFP-Booster ATTO 488 (Chromotek, GBA488-100, Lot: 90401001AT1-04). All secondary antibodies were used at a dilution of 1:1000 for immunofluorescence.

The following secondary antibodies were used for western blot experiments: IRDye 680RD Donkey anti-Guinea Pig (Li-COR, 926-68077, Lot: C70425-05); IRDye 680RD Donkey anti-Chicken (Li-COR, 926-68075, Lot: C70201-50); IRDye 800CW Donkey anti-Mouse (Li-COR, 926-32212, Lot: D10414-15); Alexa fluor 680 AffiniPure Goat anti-Mouse IgG light chain (Jackson Immuno, 115-625-174, Lot: 131312); IRDye 680RD Donkey anti-Rabbit (Li-COR, 926-68073, Lot: 91204-03, D11207-05); IRDye 800CW Donkey anti-Rabbit (Li-COR, 926-32213, Lot: D00304-05).

### Plasmids

ptf-Galectin-3 was a gift from Tamotsu Yoshimori (Addgene plasmid # 64149) (Maejima et al., 2013). GFP-Galectin-3 was subcloned from ptf-Galectin-3 (Addgene: 64149). pMXs-puro eGFP-p62 was a gift from Noboru Mizushima (Addgene plasmid # 38277) (Itakura and Mizushima, 2011). HaloTag-p62 was subcloned from eGFP-p62 (Addgene plasmid # 38277) into pHTN Halo-Tag vector (Promega). mCherry-WT-p62, mCherry-p62ΔPB1, mCherry-p62ΔUBA, mCherry-p62 K7A/D69A, and pmCherry-C1 [Clontech] were gifts from S. Martens (Max Perutz Labs, University of Vienna) (Wurzer et al., 2015). ALS-associated mutations in p62 (L341V, P392L, and G425R) were generated via site-directed mutagenesis of mCherry-WT-p62. pLJC5-Tmem192-3xHA was a gift from David Sabatini (Addgene plasmid # 102930) (Abu-Remaileh et al., 2017). pRK5-FLAG-LAMTOR2 was a gift from David Sabatini (Addgene plasmid # 42330) (Bar-Peled et al., 2012). mCherry-LAMTOR2 was subcloned from FLAG-LAMTOR2 into pmCherry-C1 [ClonTech]. eGFP-TAX1BP1 was generated by subcloning from HaloTag-TAX1BP1 (Promega) (Moore and Holzbaur, 2016). eGFP-NBR1 was a gift from Peter Kim (The Hospital for Sick Children, University of Toronto) (Deosaran et al., 2012). pAc-GFP-C1 vector was purchased from Clontech. LAMP1-KillerRed was subcloned from LAMP1-RFP. LAMP1-RFP was a gift from Walther Mothes (Addgene plasmid # 1817) (Sherer et al., 2003). LAMP1-KillerRed was subcloned into KillerRed-dMito [Evrogen]. HaloTag-WIPI2b was subcloned from GFP-WIPI2b (Dooley et al., 2014; Stavoe et al., 2019). Halo-DFCP1 was subcloned from GFP-DFCP1 (Stavoe et al., 2019). pMXs-puro GFP-DFCP1 was a gift from Noboru Mizushima (Addgene plasmid # 38269) (Itakura and Mizushima, 2010). mCherry-LC3b was subcloned from eGFP-LC3b. pEGFP-LC3 was a gift from Tamotsu Yoshimori (Addgene plasmid # 21073) (Kabeya, 2000). mCherry-LC3b was subcloned into pmCherry-C1 [Clontech]. PGK-LAMP1-mNeon was a gift from M. Ward (NIH). LAMP2-BFP was subcloned from eGFP-N1-LAMP2, which was a gift from E. Chapman (UW Madison).

### Cell culture

HeLa-M cells (referred to as HeLa) (A. Peden, Cambridge Institute for Medical Research), HEK293T, p62KO HeLa (R. Youle, NIH), and pentaKO HeLa (R. Youle, NIH), were cultured in Dulbecco’s Modified Eagle’s Medium (Corning - MT10-013-CV or Gibco - 11965084), which was supplemented with 10% Fetal Bovine Serum and 1% GlutaMax (Thermo - 35050061). Cells were cultured in a 5% CO_2_ incubator at 37 C. Cells were tested for mycoplasma contamination routinely, using MycoAlert detection kit (Lonza, LT07). Additionally, the DNA Sequencing Facility at the University of Pennsylvania authenticated the HeLa cells, using STR profiling.

Embryonic mouse neurons were dissected at E15.5, and were plated onto 0.5mg/mL poly-L-lysine coated 6-well plates for lysate collections. Primary rat hippocampal neurons were received from the Neurons R Us Culture Service Center at the University of Pennsylvania. Prior to receiving cells, 35-mm glass-bottom dishes (MatTek) were coated with 0.5mg/mL poly-L-lysine (Sigma) overnight at 37 C. Primary neurons were plated at 220,000 cells per dish into attachment media (MEM supplemented with 10% horse serum, 33mM D-glucose, and 1mM sodium pyruvate). Following attachment (4-6hrs), attachment media was removed and replaced with maintenance media (Neurobasal [Gibco] supplemented with 33mM D-glucose [Sigma], 2mM GlutaMax [Gibco], Penicilin (100units/mL) / Streptomycin (100µg/mL) [Giibco], and 2% B27 [Gibco]). On the following day, 5µM AraC [Sigma] was added to the cultures to prevent proliferation of non-neuronal cells. Neurons were cultured with 5% CO_2_ at 37 C for 7-10 days prior to transfection.

Human LAMP1-eGFP i^3^Neuron iPSCs were generated previously (Boecker et al., 2020). iPSCs were cultured in Essential 8 medium [ThermoFisher] and plated onto Matrigel coated dishes. An established protocol was used for neuronal differentiation into i^3^Neurons (Fernandopulle et al., 2018). Following differentiation, i^3^Neurons were cryo-preserved in i^3^Neuron media (BrainPhys Neuronal Medium supplemented [StemCell] with 2% B27 [Gibco], 10ng/mL BDNF [PreproTech], 10ng/mL NT-3 [PeproTech], and 1µg/mL Laminin [Corning]) with 10% DMSO added. Prior to culturing i^3^Neurons, 35-mm glass-bottom dishes [MatTek] were coated with poly-L-ornithine overnight at 37 C. i^3^Neurons were cultured for 21-22 days in 5% CO_2_ at 37 C and were fed every 3-4 days with fresh culture media.

### Transfection

For fixed and live-cell experiments without siRNA, HeLa cells, p62KO HeLa cells, and pentaKO cells were transfected with 0.3-1.5μg of DNA using FuGene 6 [Promega] and incubated for 24hrs. However, HeLa cells transfected with LAMP1-KillerRed were incubated for 48hrs.

In knockdown experiments, HeLa cells were transfected with 40µM siRNA using Lipofectamine RNAiMax and incubated for 48hrs. To knockdown p62, ON-TARGETplus p62 siRNA (J-010230-05) was used on HeLa cells (Target sequence: ‘GAACAGAUGGUCGGAUA’, Horizon Discovery). To knockdown HSP27, ON-TARGETplus HSPB1 pool siRNA was using on HeLa cells (Target sequences: 1: ‘CAAGUUUCCUCCUCCCUGU,’ 2: ‘GAGACUGCCGCCAAGUAAA,’ 3: ‘GGUGCUUCACGCGGAAAUA,’ 4: ‘CCACGCAGUCCAACGAGAU,’ Horizon Discovery). In knockdown and rescue experiments, HeLa cells were transfected with 40µM siRNA alongside 0.5-1.0µg total DNA using Lipofectamine 2000 and incubated for 48hrs.

Rat hippocampal neurons were transfected following 8-11 days *in vitro*. Neurons were transfected with 0.5-1.0µg of DNA using Lipofectamine 2000. Neurons were incubated with DNA:Lipid complexes for 45 min prior to replacement with full conditioned media. Neurons were fixed for immunofluorescence 24hrs later. However, rat hippocampal neurons transfected with LAMP1-KillerRed were incubated for 48hrs prior to live-cell imaging.

### LLOMe-induced lysosomal damage

L-Leucyl-L-Leucine methyl ester monohydrochloride (Cayman) was dissolved in 100% ethanol. In fixed cell experiments, HeLa, p62KO, PentaKO were treated with 250µM or 750µM LLOMe for 2 hr prior to fixation. In live cell experiments, HeLa cells were removed from culture media and replaced with HeLa imaging media (Leibovitz’s L-15 Medium (Gibco) supplemented with 10% FBS and 1% GlutaMax). Subsequently, 750µM LLOMe was added to HeLa imaging media. For immunofluorescence experiments, rat hippocampal neurons and i^3^Neurons were treated with 1.0mM LLOMe for 2 hr prior to fixation.

### Stable cell line generation

Stable cell lines were generated using lentivirus transduction and subsequent single-cell isolation. In brief, 700,000 HEK293T cells were seeded in a 6-cm dish. The following day, HEK293T cells were transfected with 1µg (LAMP1-mNeonGreen) or 3µg (TMEM192-3xHA) along with packaging plasmids (psPAX2 and Vsv-G) using Fugene 6 [Promega]. HEK293T cells were incubated for either 24hr (LAMP1-mNeonGreen) or 36hr (TMEM192-3xHA). Following this, the media was replaced with complete culture media supplemented with Viral Boost Reagent [ALSTEM]. After 24 hours, the conditioned media was collected for HeLa transduction and passed through a 45um filter.

HeLa were seeded at 50,000 cells in a 6-cm dish on the day before transduction. HeLa media was removed and replaced with virus for 24hrs. Following the transduction, cells were grown until 70% confluency and then single cells were isolated and plated onto a 96-well plate. To isolate single cells, cells were diluted to a final concentration of 5 cells/mL, using an established protocol [Addgene]. Clones were selected by expression level of construct of interest.

### Immunoprecipitation

Lysosomal immunoprecipitation experiments were performed following an established protocol (Abu-Remaileh et al., 2017). In brief, HeLa TMEM192-3xHA were seeded at 4 million cells/15-cm dish. Approximately 48hr later when cells reach 80% confluence, cells were treated with 1.0mM LLOMe for 1hr or were untreated. Cells were scraped into cold KPBS buffer (1x PBS (50mM NaPO_4_, 150mM NaCl, pH=7.4) supplemented with 136mM KCl, 10mM KH_2_PO_4_, 50mM sucrose, 1mM Phenylmethanesulfonyl Fluoride, 20µg/mL TAME, 20µg/mL Leupeptin, 2µg/mL Pepstatin-A). In the absence of sucrose, LLOMe-treated lysosomes had consistently decreased yields. Cells were spun down and resuspended into 950uL from which 50µL was stored on ice as whole cell fraction. The remainder of the cell suspension was homogenized using 2mL glass homogenizer. Following this, the homogenized lysate is spun down, collecting the resultant supernatant as lysate fraction. Protein concentration of lysate was quantified using a Bradford Concentration Assay (BCA), and then protein concentration was normalized across conditions. Following normalization, 100µL of anti-HA magnetic beads were added to the remainder of the sample. Following incubation, magnetic beads were washed with KPBS buffer containing 300mM NaCl prior to elution. Magnetic beads were eluted into 80µL SDS sample buffer.

eGFP-HSP27 immunoprecipitations were performed in p62KO HeLa cells 24hrs following transient transfection using Fugene 6 (Promega). HeLa cells were treated with 750µM LLOMe for 2hrs prior to immunoprecipitation. Cells were collected into and incubated in lysis buffer (ddH_2_0 supplemented with 20mM Tris-HCl, 137mM NaCl, 2mM EDTA, 1% NP-40, 5% Glycerol, 1mM Phenylmethanesulfonyl Fluoride, 20µg/mL TAME, 20µg/mL Leupeptin, 2µg/mL Pepstatin-A). Following 15min of lysis, lysates were spun, and the supernatant was collected as the lysate fraction. 600µL of lysate was added to Protein G DynaBeads, which had been incubated with 1.25µg Mouse anti-GFP (Abcam, ab1218) for 30 minutes at 4 C. The magnetic beads were then washed with TBS supplemented with 0.05% Tween-20 (0.05% TBS-T). A small fraction of lysate was stored on ice for lysate fraction. Dynabeads were incubated in lysate for 1hr at 4 C, and then were washed with 0.05% TBS-T prior to elution. Magnetic beads were eluted into 60µL SDS sample buffer.

### Western blotting

To assay expression levels of proteins in HeLa, i^3^Neurons, and mouse cortical neurons, cells were washed 2x with ice cold PBS prior to lysis with RIPA buffer (50mM Tris-HCl, 150mM NaCl, 0.1% Triton X-100, 0.5% deoxycholate, 0.1% SDS, 2x Halt Protease and Phosphatase inhibitor). Cells were snap frozen in liquid nitrogen prior to incubation in RIPA buffer for 30 min at 4 C. Samples were then centrifuged, and the supernatant was collected as the lysate fraction. A BCA was performed on the collected lysate, and samples were denatured in sample buffer containing SDS at 95 C.

For immunoprecipitation experiments and protein expression experiments, samples were resolved using SDS-PAGE gels. Following electrophoresis, proteins were transferred to Immobilon-FL PVDF membranes [Millipore]. The membrane was dried for one hour prior to rehydration in methanol. The membranes were then stained for total proteins, using Li-COR Revert Total Protein stain. The membranes were imaged using an Odyssey CLx Infrared Imaging System (Li-COR). Following imaging, membranes were destained with 0.1M NaOH supplemented with 30% Methanol.

Membranes were then blocked in either TrueBlack WB Blocking Buffer (Biotium) or EveryBlot Blocking Buffer (Bio-Rad). When using TrueBlack WB Blocking Buffer, membranes were blocked for one hour and were incubated with primary antibodies diluted in TrueBlack WB Antibody Diluent (Biotium) supplemented with 0.2% Tween-20 (Bio-Rad) at 4 C overnight. Alternatively, when membranes were blocked with EveryBlot Blocking Buffer, membranes were blocked for 5-10 minutes and were incubated with primary antibodies diluted in EveryBlot Blocking Buffer overnight at 4 C. After 12-16 hours incubation, membranes were washed 3x with 1xTBS (50mM Tris-HCl, 274mM NaCl, 9mM KCl) supplemented with 0.1% Tween-20 (TBS-T). Membranes were then incubated in secondary antibody, which were diluted in either TrueBlack WB Blocking Buffer supplemented with 0.01% SDS or EveryBlot Blocking Buffer supplemented with 0.02% SDS. Membranes were incubated for one hour and then were washed 3x with TBS-T. Membranes were then imaged, and band intensities were measured in the Li-COR Image Studio application.

### Fixation and permeabilization conditions

A variety of fixation methods were used, depending on antibodies and cell types used and following optimization. In experiments assaying LC3b intensity in HeLa, PentaKO, or p62KO cells, cells were fixed using Bouin’s Solution supplemented with 8% sucrose. Bouin’s solution was added in a 1:1 ratio with cell media, and cells were incubated for 30min at room temperature. Cells were washed with PBS prior to permeabilization with ice-cold methanol. In experiments assaying the effect of ALS-associated mutants on LC3b intensity in hippocampal neurons or in HeLa cells, cells were fixed in ice-cold methanol.

In experiments assaying WIPI2b-puncta number, cells were fixed with 4% paraformaldehyde supplemented with 4% sucrose (4% PFA/4% sucrose). Cells were incubated with Janelia Fluor 646-Halo ligand [Promega] for 1.5 hrs in blocking solution (5% goat serum and 1% BSA in PBS).

In experiments assaying co-localization of LAMP1 with receptors, with GABARAP, or with eGFP-HSP27 in i^3^Neurons or in HeLa cells, cells were fixed with 4% PFA/4% sucrose. Permeabilization was performed with ice-cold methanol. p62 localization experiments in p62KO and pentaKO cells as well as in hippocampal neurons were fixed with 4% PFA/4% sucrose. In the HeLa cells permeabilization was performed with ice-cold methanol. The hippocampal neurons were mounted in ProLong Gold anti-fade mountant (ThermoFisher). Similarly, in eGFP-HSP27 puncta formation assays in i^3^Neurons or p62KO cells, cells were fixed with 4% PFA/4% sucrose before mounting in ProLong Gold anti-fade mountant.

### Immunostaining

Following permeabilization, cells were blocked in blocking solution (5% goat serum and 1% BSA in PBS) for 90 min. Samples were then incubated with primary antibodies overnight at 4 C. Primary antibodies were diluted into blocking solution. After 14-16 hours of incubation, cells were washed with PBS and then incubated in secondary antibodies. Secondary antibodies were also diluted into blocking solution. Following secondary antibody incubations, samples were washed with PBS. Frequently, samples were treated with a nuclear counterstain [Hoechst 33342, Invitrogen] for 10 min. Coverslips were mounted in ProLong Gold [Life Technologies].

Images were acquired using a Perkin Elmer UltraView Vox spinning disk confocal on a Nikon Eclipse Ti Microscope. Fixed cell experiments were performed on an Apochromat 100x 1.49NA oil-immersion objective, and live-cell experiments were performed on a Plan Aprochromat Lambda 60x 1.40NA oil-immersion objective. Z-stacks were collected at 200nm step-size. Experiments were imaged on either a Hamamatsu EMCCD C9100-50 camera or a Hamamatsu CMOS ORCA-Fusion (C11440-20UP). The EMCCD camera was used with Volocity Software [Quorom Technologies/PerkinElmer]. The CMOS camera was used with VisiView (Visitron).

### Immunofluorescence quantifications

To measure LC3b/GABARAP intensities at either Flag-LAMTOR2 or mCherry-LAMTOR2 puncta, max projections of each channel in each image were made. Individual cells were selected from these max projections. Cells were selected if they were completely in frame of the image, were equivalently transfected, were in interphase, and were not blebbing or going through apoptosis. In these cells, LAMTOR2 area was thresholded and binarized using Otsu thresholding in ImageJ (Otsu, 1979). The binarized image was used to generate regions of interest, and then LC3b/GABARAP fluorescence intensity was measured within these regions of interest.

To quantify area within the cell occupied by an individual protein, we generated max projections of each channel in every image. Individual cells were selected from these max projections, using similar parameters as listed above. We then segmented the area of these proteins using Ilastik, a machine-learning based approach to image segmentation (Berg et al., 2019). The percentage overlap between different proteins was measured using the image calculator function in ImageJ.

### Live-Cell Imaging

For live-cell imaging of HeLa cells, the culture media was replaced with Leibovitz’s L-15 media (Gibco, 11415064). The media was supplemented with 1% Glutamax and 10% fetal bovine serum. Cells were then moved onto the microscope stage, which is surrounded by a 37 C imaging chamber. The samples were given several minutes to equilibrate before imaging. When using LLOMe, a single frame was captured prior to lysosomal damage treatment. LLOMe was then added directly to the imaging dish, and then cells were imaged for 1 or 2 hours. To avoid photo toxicity, cells were imaged a low frame rate (ranging from one frame every minute to one frame every ten minutes), depending on the experiment.

In KillerRed experiments, both HeLa and rat hippocampal neurons underwent 7 photo-bleaching cycles with 9ms per pixel. Cells were imaged for either 45 minutes or 1 hr post-initial FRAP cycle.

In experiments using live-cell imaging of rat hippocampal neurons, culture media was removed and replaced with Hibernate E medium, which was supplemented with 2% B27 and 33mM D-glucose.

## Acknowledgements

We thank Mariko Tokito for technical assistance, and we thank Adam Fenton, Alex Boecker, Sierra Palumbos, and Juliet Goldsmith for insight and discussion. This work was supported by National Institutes of Health Grants T32 GM007229 and F31 NS125954-01 to E.R.G, and R01 NS060698 to E.L.F.H.

## Contributions

E.R.G and E.L.F.H devised experiments and wrote the manuscript. E.R.G performed the experiments and subsequent analysis with supervision by E.L.F.H.

## Resource Availability

### Materials availability

All unique reagents generated in this study are available from the Lead Contact with a completed Materials Transfer Agreement.

### Lead Contact

For requests regarding reagents should be directed to and will be fulfilled by the Lead Contact, E.L.F.H.

