## Supplemental Figures for "The selective autophagy receptor p62 and the heat shock protein HSP27 facilitate lysophagy via the formation of phase-separated condensates"

Supplemental Video 1 – **Activation of LAMP1-KillerRed induces recruitment of p62 to lysosomes in HeLa cells** [Related to Figure 1].

Representative video demonstrating recruitment of p62 to lysosomes following activation of LAMP1-KillerRed with irradiation with 561nm light. Prior to photo-bleaching, LAMP1-KillerRed (cyan) co-localizes with LAMP2-BFP (magenta) (first panel). Approximately ten minutes following photo-bleaching, eGFP-p62 (green) begins to accumulate on LAMP2-BFP vesicles (second panel). Accumulation of p62 is visible in single greyscale channel (third panel). Scale bar: 10μm. Stopwatch shows time in minutes:seconds.

Supplemental Video 2 – **Activation of LAMP1-KillerRed induces recruitment of p62 to lysosomes in primary hippocampal neurons** [Related to Figure 2].

Representative video demonstrating p62 recruitment to damaged lysosomes following LAMP1-KillerRed activation in primary rat hippocampal neurons, following 561nm light irradiation. Prior to photo-bleaching, LAMP1-KillerRed (first panel) co-localizes with LAMP2-BFP (second panel) (magenta). Approximately five minutes following photo-bleaching, eGFP-p62 (third panel) (green) begins to accumulate on LAMP2-BFP vesicles (fourth panel). Scale bar: 2μm. Stopwatch shows time in minutes:seconds.

Supplemental Video 3 – **Addition of LLOMe results in LAMTOR2 and galectin-3 localization to lysosomes** [Related to Figure 3].

Representative video demonstrating that the addition of 750μM LLOMe results in localization of both LAMTOR2 and galectin-3 to damaged lysosomes in HeLa cells. In the top row, HeLa cells were treated with ethanol and imaged for one hour. In the bottom row, HeLa cells were treated with LLOMe. Both LAMTOR2 and galectin-3 form puncta rapidly, as shown in single channel images for galectin-3 (second column, green) and for LAMTOR (third column, magenta). LAMTOR2 and galectin-3 co-localization with LAMP1-Halo (blue) is observed in the merged image (first column). Scale bar: 10μm. Time 0 indicates when LLOMe was added. Stopwatch shows time in minutes:seconds.

Supplemental Video 4 – **HSP27 is recruited to damaged lysosomes** [Related to Figure 5].

Representative video demonstrating that HSP27 is recruited to LAMTOR2 puncta following the addition of 750μM LLOMe in HeLa cells. LAMTOR2 forms puncta following LLOMe addition (third column, magenta), and HSP27 is recruited to these puncta after approximately forty minutes of LLOMe treatment (second column, green). Arrowheads indicate example co-localization events. Scale bar: 2μm. Time 0 indicates when LLOMe was added. Stopwatch shows time in minutes:seconds.

Supplemental Video 5 – **HSP27 co-localizes with Double FYVE-containing protein 1 (DFCP1) following lysosomal damage** [Related to Figure 5].

Representative video demonstrating that HSP27 co-localizes with DFCP1 following the addition of 750μM LLOMe in HeLa cells. DFCP1 forms puncta following LLOMe addition (second column, magenta), and HSP27 co-localizes both spatially and temporally with DFCP1 (first column, green). Arrowheads indicate example co-localization events. Scale bar: 2μm. Time 0 indicates when LLOMe was added. Stopwatch shows time in minutes:seconds

Supplemental Video 6 – **HSP27 is recruited to p62 puncta following lysosomal damage** [Related to Figure 6].

Representative video demonstrating the HSP27 is recruited to p62 following the addition of 750μM LLOMe in HeLa cells. p62 puncta are observed in single channel images following the addition of LLOMe (first column, green). HSP27 is recruited to p62 following the addition of LLOMe (second column, magenta). Arrowheads indicate example co-localization events. Scale bar: 2μm. Time 0 indicates when LLOMe was added. Stopwatch shows time in minutes:seconds

Supplemental Video 7 – **Depletion of HSP27 decrease the liquidity of p62 condensates** [Related to Figure 6].

Representative video demonstrating that HeLa cells transiently transfected with HSP27 siRNA have decreased fluorescence recovery of p62 condensates. In the top row, HeLa cells were transfected with control siRNA. In the bottom row, HeLa cells were transfected with HSP27 siRNA. In the merged image, eGFP-p62 (green, second column) co-localizes with LAMP1-RFP (magenta) prior to photo-bleaching. Arrowheads indicate the location of eGFP-p62 rings following photo-bleaching. HeLa cells were treated with 750μM LLOMe for one hour prior to imaging. Cells were imaged for one minute prior to photo-bleaching. Stopwatch shows time in minutes:seconds

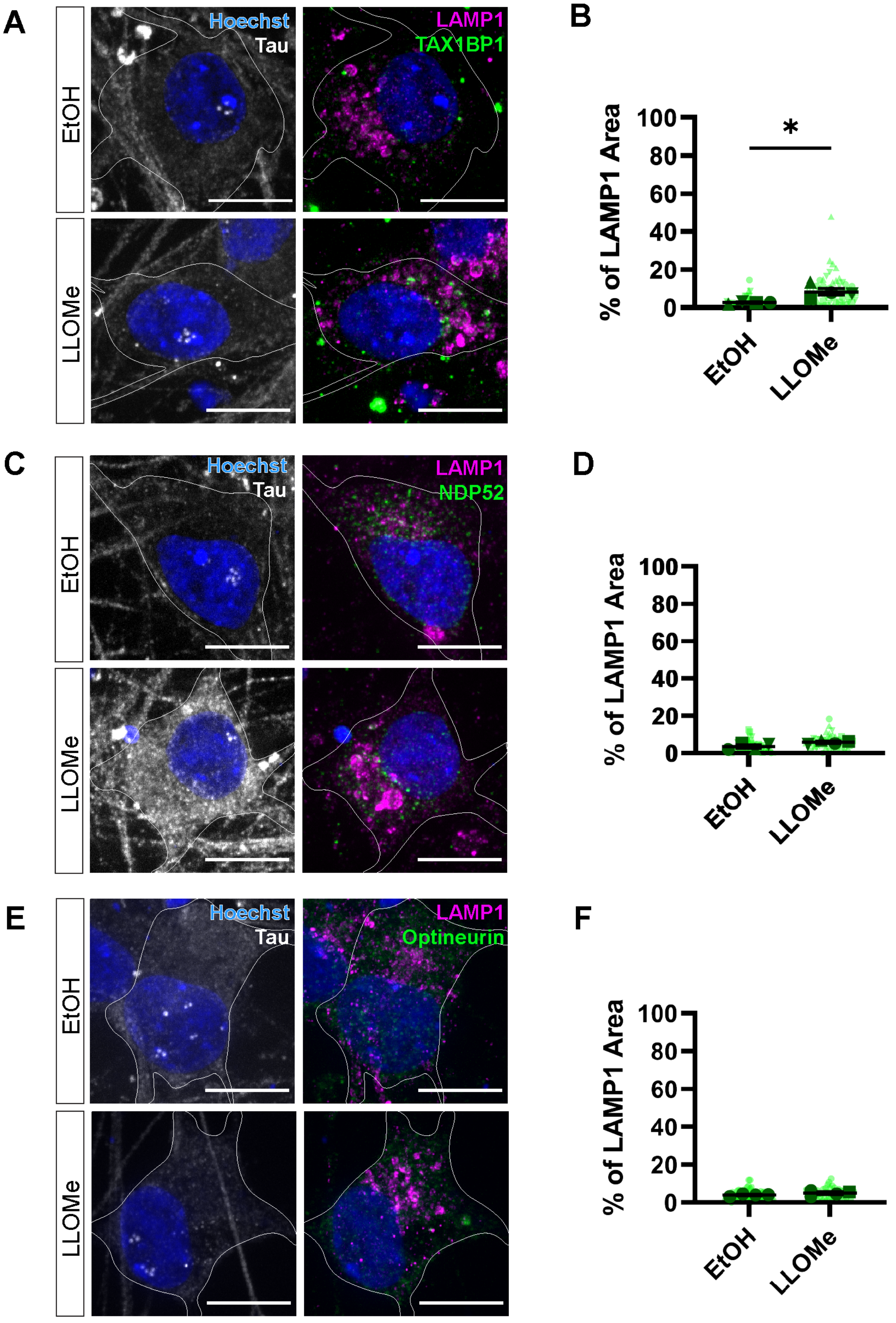


Supplementary Figure 1 – **Damaged lysosomes recruit the selective autophagy receptor TAX1BP1 in i³Neurons** [Related to Figure 2].

(A) Representative images of anti-TAX1BP1 localization to lysosomes in i³Neurons following treatment with either EtOH or 1.0mM LLOMe for two hours. Cell outlines were traced. Scale Bar: 10µm.

(B) Quantification of the percentage of LAMP1 area occupied by anti-TAX1BP1 across conditions (Unpaired t-test, $p < 0.05$, N=4, $n_{(EtOH)}$= 45, $n_{(LLOMe)}$=56).

(C) Representative images of anti-NDP52 localization to lysosomes in i$^3$Neurons following treatment with either EtOH or 1.0mM LLOMe for two hours. Cell outlines were traced. Scale Bar: 10µm.

(D) Quantification of the percentage of LAMP1 area occupied by anti-NDP52 across conditions (Unpaired t-test, p = 0.0708, N=4, $n_{(EtOH)}$= 59, $n_{(LLOMe)}$=45).

(E) Representative images of anti-Optineurin localization to lysosomes in i$^3$Neurons following treatment with either EtOH or 1.0mM LLOMe for two hours. Cell outlines were traced. Scale Bar: 10µm.

(F) Quantification of the percentage of LAMP1 area occupied by anti-Optineurin across conditions (Unpaired t-test, p = 0.2237, N=4, $n_{(EtOH)}$= 59, $n_{(LLOMe)}$=50).

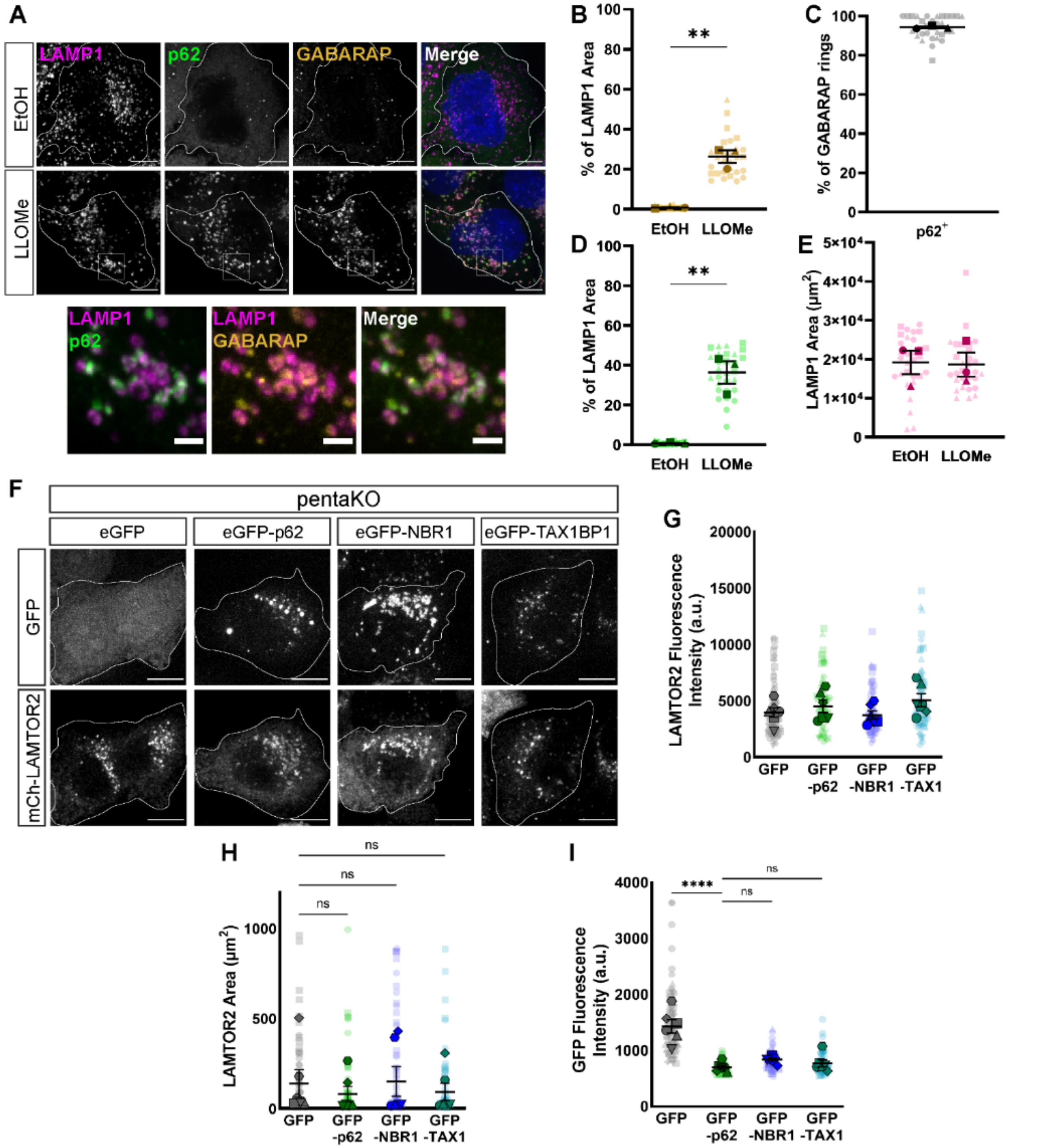


Supplemental Figure 2 – **Expression of p62 is sufficient to drive lysophagy in pentaKO HeLa cells** [Related to Figure 4].

(A) Representative max projections demonstrating the co-localization of anti-LAMP1, anti-p62, anti-GABARAP in HeLa cells, which were treated with EtOH or 750µM LLOMe for 2 hours. Cell outlines are traced. White boxes indicate inset image region. Scale Bar: 10µm; inset: 2µm.

(B) Quantification of the percentage of LAMP1 occupied by anti-GABARAP (Unpaired t-test, $p < 0.01$, N=3, $n_{EtOH} = 30$, $n_{LLOMe} = 29$).
(C) Quantification of the percentage of anti-GABARAP rings (in single z-plane) co-localizing with anti-p62.
(D) Quantification of the percentage of LAMP1 occupied by anti-p62 (Unpaired t-test, $p < 0.01$, N=3, $n_{EtOH} = 30$, $n_{LLOMe} = 29$).
(E) Quantification of LAMP1 area per cell (Unpaired t-test, $p = 0.9117$, N=3, $n_{EtOH} = 30$, $n_{LLOMe} = 29$).
(F) Representative images of GFP and mCherry-LAMTOR2 in pentaKO HeLa cells. Cells were treated with 750µM LLOMe for two hours. Cells were transfected with mCherry-LAMTOR2 and either GFP vector, eGFP-p62, eGFP-NBR1, or eGFP-TAX1BP1. Cell outlines are traced. Scale Bar: 10µm.
(G) Quantification of mCherry-LAMTOR2 fluorescence intensity across the conditions (One-way ANOVA, $p_{(GFP/p62)} = 0.7510$, $p_{(GFP/NBR1)} = 0.9785$, $p_{(GFP/TAX1)} = 0.2771$, N=6, $n_{GFP} = 72$, $n_{p62} = 64$, $n_{NBR1} = 77$, $n_{TAX1} = 74$).
(H) Quantification of mCherry-LAMTOR2 area across the conditions (One-way ANOVA, $p_{(GFP/p62)} = 0.8614$, $p_{(GFP/NBR1)} = 0.9986$, $p_{(GFP/TAX1)} = 0.9188$, N=6, $n_{GFP} = 72$, $n_{p62} = 64$, $n_{NBR1} = 77$, $n_{TAX1} = 74$).
(I) Quantification of GFP fluorescence intensity across the conditions (One-way ANOVA, $p_{(p62/GFP)} < 0.0001$, $p_{(p62/NBR1)} = 0.3804$, $p_{(p62/TAX1)} = 0.7963$, N=6, $n_{GFP} = 72$, $n_{p62} = 64$, $n_{NBR1} = 77$, $n_{TAX1} = 74$).

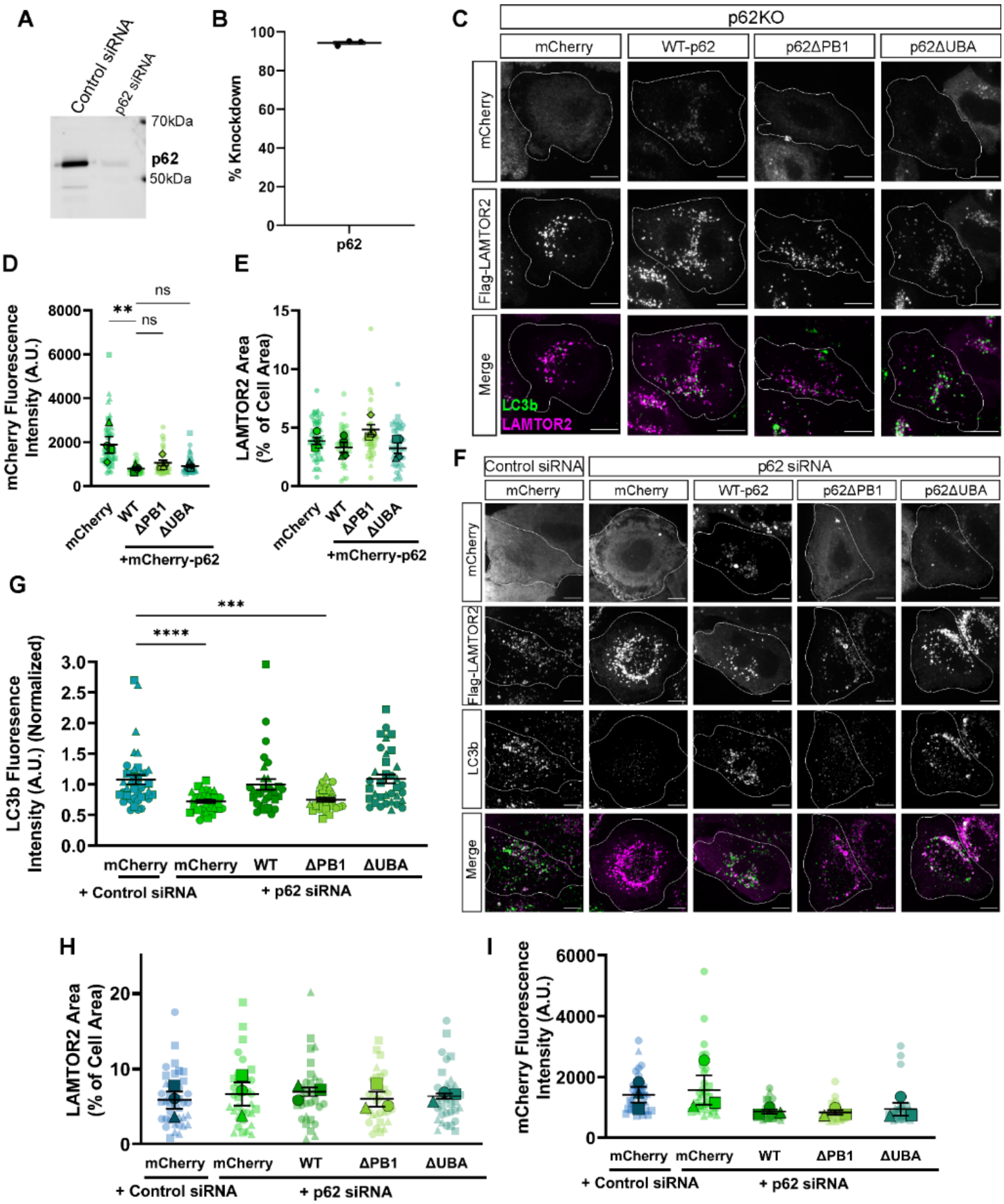


Supplemental Figure 3 – **p62 is required for lysophagy** [Related to Figure 4].

(A) Western blot analysis of cell lysates from HeLa cells, evaluating expression of p62 following transient transfection of either control siRNA or p62 siRNA.

(B) Quantification of percentage knockdown in HeLa cells that were transfected with p62 siRNA relative to total protein and to control siRNA (94 ± 0.77% knock down).

(C) Representative images of anti-LC3b, mCherry, and Flag-LAMTOR2 in p62KO HeLa cells. Cells were transiently transfected with Flag-LAMTOR2 and either mCherry vector, WT-p62, p62ΔPB1, or p62ΔUBA. Cells were treated with LLOMe for two hours. Cell outlines are traced. Scale Bar: 10µm.

(D) Quantification of mCherry fluorescence intensity per cell across different conditions (One-way ANOVA, $p_{(WT/mCherry)} < 0.01$, $p_{(WT/\Delta PB1)} = 0.6323$, $p_{(WT/\Delta UBA)} = 0.6933$, N=4, $n_{mCherry} = 44$, $n_{WT} = 36$, $n_{\Delta PB1} = 39$, $n_{\Delta UBA} = 44$).

(E) Quantification of Flag-LAMTOR2 area per cell across different conditions (One-way ANOVA, $p_{(WT/mCherry)} = 0.5887$, $p_{(WT/\Delta PB1)} = 0.0635$, $p_{(WT/\Delta UBA)} = 0.9073$, N=4, $n_{mCherry} = 44$, $n_{WT} = 36$, $n_{\Delta PB1} = 39$, $n_{\Delta UBA} = 44$).

(F) Representative images of HeLa cells that were transiently transfected with either control siRNA or p62 siRNA. Cells were co-transfected with Flag-LAMTOR2 and either mCherry vector, WT-p62, p62ΔPB1, or p62ΔUBA. Cell outlines are traced. Scale Bar: 10µm.

(G) Quantification of anti-LC3b intensity at Flag-LAMTOR2 area (Kruskal-Wallis test, $p_{(control\ si\ +\ mCherry/p62\ si\ +\ mCherry)} < 0.0001$, $p_{(control\ si\ +\ mCherry/p62\ si\ +\ WT)} = 0.8909$, $p_{(control\ si\ +\ mCherry/p62\ si\ +\ \Delta PB1)} < 0.001$, $p_{(control\ si\ +\ mCherry/p62\ si\ +\ \Delta UBA)} > 0.9999$, $N_{control\ si\ +\ mCherry} = 39$, $N_{p62\ si\ +\ mCherry} = 34$, $N_{p62\ si\ +\ WT} = 31$, $N_{p62\ si\ +\ \Delta PB1} = 30$, $N_{p62\ si\ +\ \Delta UBA} = 35$).

(H) Quantification of Flag-LAMTOR2 area per cell across different conditions (One-way ANOVA, $p_{(control\ si\ +\ mCherry/p62\ si\ +\ mCherry)} = 0.9247$, $p_{(control\ si\ +\ mCherry/p62\ si\ +\ WT)} = 0.9150$, $p_{(control\ si\ +\ mCherry/p62\ si\ +\ \Delta PB1)} = 0.9290$, $p_{(control\ si\ +\ mCherry/p62\ si\ +\ \Delta UBA)} = 0.9247$, N=3, $n_{control\ si\ +\ mCherry} = 39$, $n_{p62\ si\ +\ mCherry} = 34$, $n_{p62\ si\ +\ WT} = 31$, $n_{p62\ si\ +\ \Delta PB1} = 30$, $n_{p62\ si\ +\ \Delta UBA} = 35$).

(I) Quantification of mCherry fluorescence intensity across different conditions (One-way ANOVA, $p_{(control\ si\ +\ mCherry/p62\ si\ +\ mCherry)} = 0.6894$, $p_{(control\ si\ +\ mCherry/p62\ si\ +\ WT)} = 0.4862$, $p_{(control\ si\ +\ mCherry/p62\ si\ +\ \Delta PB1)} = 0.4862$, $p_{(control\ si\ +\ mCherry/p62\ si\ +\ \Delta UBA)} = 0.4862$, N=3, $n_{control\ si\ +\ mCherry} = 39$, $n_{p62\ si\ +\ mCherry} = 34$, $n_{p62\ si\ +\ WT} = 31$, $n_{p62\ si\ +\ \Delta PB1} = 30$, $n_{p62\ si\ +\ \Delta UBA} = 35$).

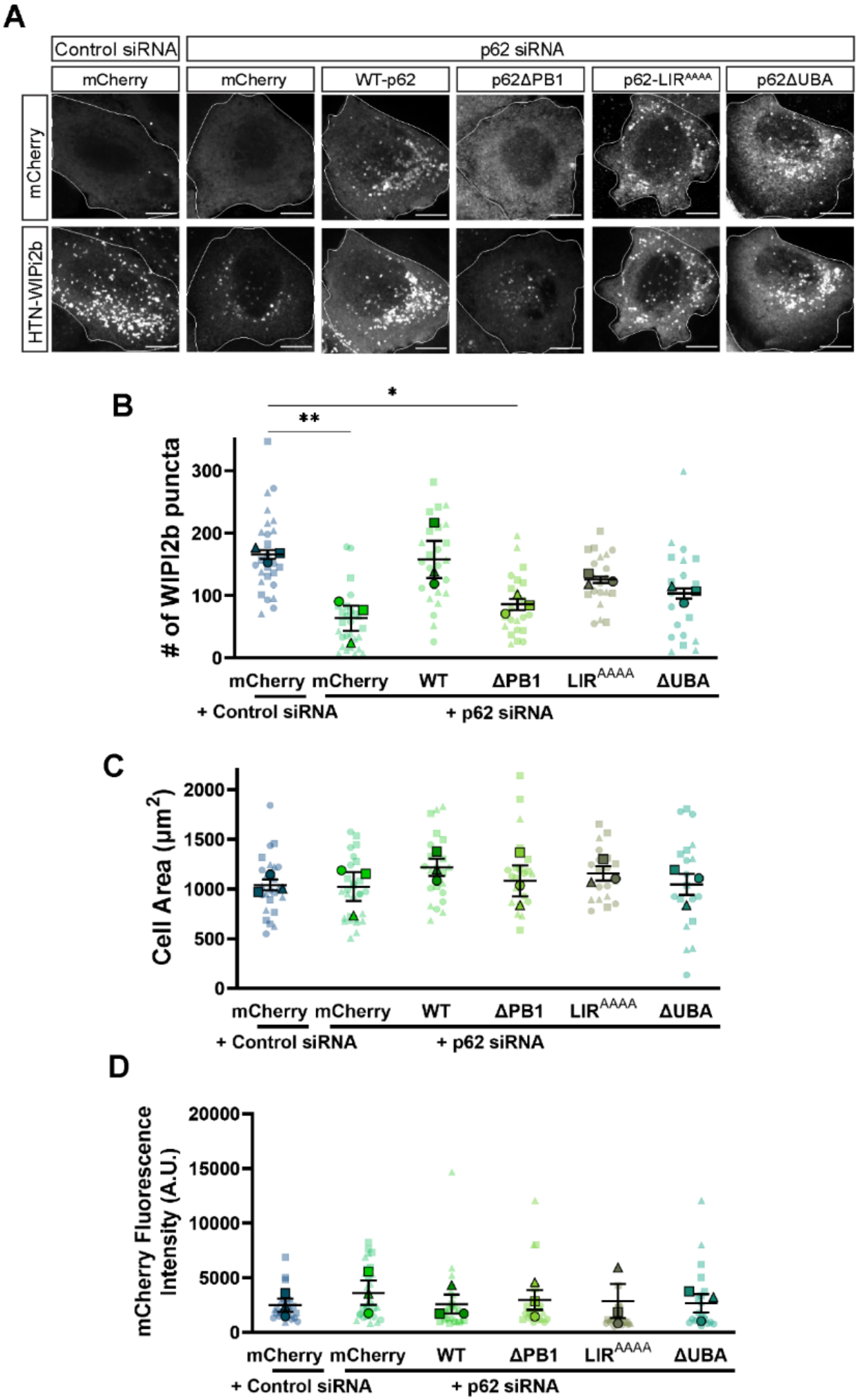
A
Control siRNA
p62 siRNA
mCherry
mCherry
WT-p62
p62ΔPB1
p62-LIR^AAAA
p62ΔUBA
mCherry
HTN-WIPI2b
B
### of WIPI2b puncta
mCherry mCherry WT ΔPB1 LIR^AAAA ΔUBA
+ Control siRNA
+ p62 siRNA
C
Cell Area (μm²)
mCherry mCherry WT ΔPB1 LIR^AAAA ΔUBA
+ Control siRNA
+ p62 siRNA
D
mCherry Fluorescence Intensity (A.U.)
mCherry mCherry WT ΔPB1 LIR^AAAA ΔUBA
+ Control siRNA
+ p62 siRNA

Supplemental Figure 4 – **p62 is crucial for induction of WIPI2b puncta in lysophagy** [Related to Figure 4].
(A) Representative images of mCherry and HaloTag-WIPI2b in HeLa cells, which were co-transfected with control siRNA or p62 siRNA. Cells were transfected with mCherry Vector, WT-p62, p62ΔPB1, p62-LIR$^{AAAA}$, or p62ΔUBA. Cells were treated with 750μM LLOMe for two hours. Cell outlines are traced. Scale Bar: 10μm.
(B) Quantification of the number of WIPI2b puncta per cell following LLOMe treatment (One-way ANOVA, $p_{(control\ si\ +\ mCherry/p62\ si\ +\ mCherry)} < 0.01$, $p_{(control\ si\ +\ mCherry/p62\ si\ +\ WT)} = 0.9955$, $p_{(control\ si\ +\ mCherry/p62\ si\ +\ \Delta PB1)} < 0.05$, $p_{(control\ si\ +\ mCherry/p62\ si\ +\ LIRmut)} = 0.3075$, $p_{(control\ si\ +\ mCherry/p62\ si\ +\ \Delta UBA)} = 0.0623$, N=3, $n_{control\ si\ +\ mCherry} = 26$, $n_{p62\ si\ +\ mCherry} = 28$, $n_{p62\ si\ +\ WT} = 24$, $n_{p62\ si\ +\ \Delta PB1} = 22$, $n_{p62\ si\ +\ LIRmut} = 20$, $n_{p62\ si\ +\ \Delta UBA} = 20$).
(C) Quantification of cell area per cell across the conditions (One-way ANOVA, $p_{(control\ si\ +\ mCherry/p62\ si\ +\ mCherry)} = 0.9999$, $p_{(control\ si\ +\ mCherry/p62\ si\ +\ WT)} = 0.6771$, $p_{(control\ si\ +\ mCherry/p62\ si\ +\ \Delta PB1)} = 0.9985$, $p_{(control\ si\ +\ mCherry/p62\ si\ +\ LIRmut)} = 0.9029$, $p_{(control\ si\ +\ mCherry/p62\ si\ +\ \Delta UBA)} > 0.9999$, N=3, $n_{control\ si\ +\ mCherry} = 26$, $n_{p62\ si\ +\ mCherry} = 28$, $n_{p62\ si\ +\ WT} = 24$, $n_{p62\ si\ +\ \Delta PB1} = 22$, $n_{p62\ si\ +\ LIRmut} = 20$, $n_{p62\ si\ +\ \Delta UBA} = 20$).
(D) Quantification of mCherry fluorescence intensity across the conditions (One-way ANOVA, $p_{(control\ si\ +\ mCherry/p62\ si\ +\ mCherry)} = 0.8908$, $p_{(control\ si\ +\ mCherry/p62\ si\ +\ WT)} > 0.9999$, $p_{(control\ si\ +\ mCherry/p62\ si\ +\ \Delta PB1)} = 0.9965$, $p_{(control\ si\ +\ mCherry/p62\ si\ +\ LIRmut)} = 0.9985$, $p_{(control\ si\ +\ mCherry/p62\ si\ +\ \Delta UBA)} = 0.9998$, N=3, $n_{control\ si\ +\ mCherry} = 26$, $n_{p62\ si\ +\ mCherry} = 28$, $n_{p62\ si\ +\ WT} = 24$, $n_{p62\ si\ +\ \Delta PB1} = 22$, $n_{p62\ si\ +\ LIRmut} = 20$, $n_{p62\ si\ +\ \Delta UBA} = 20$).

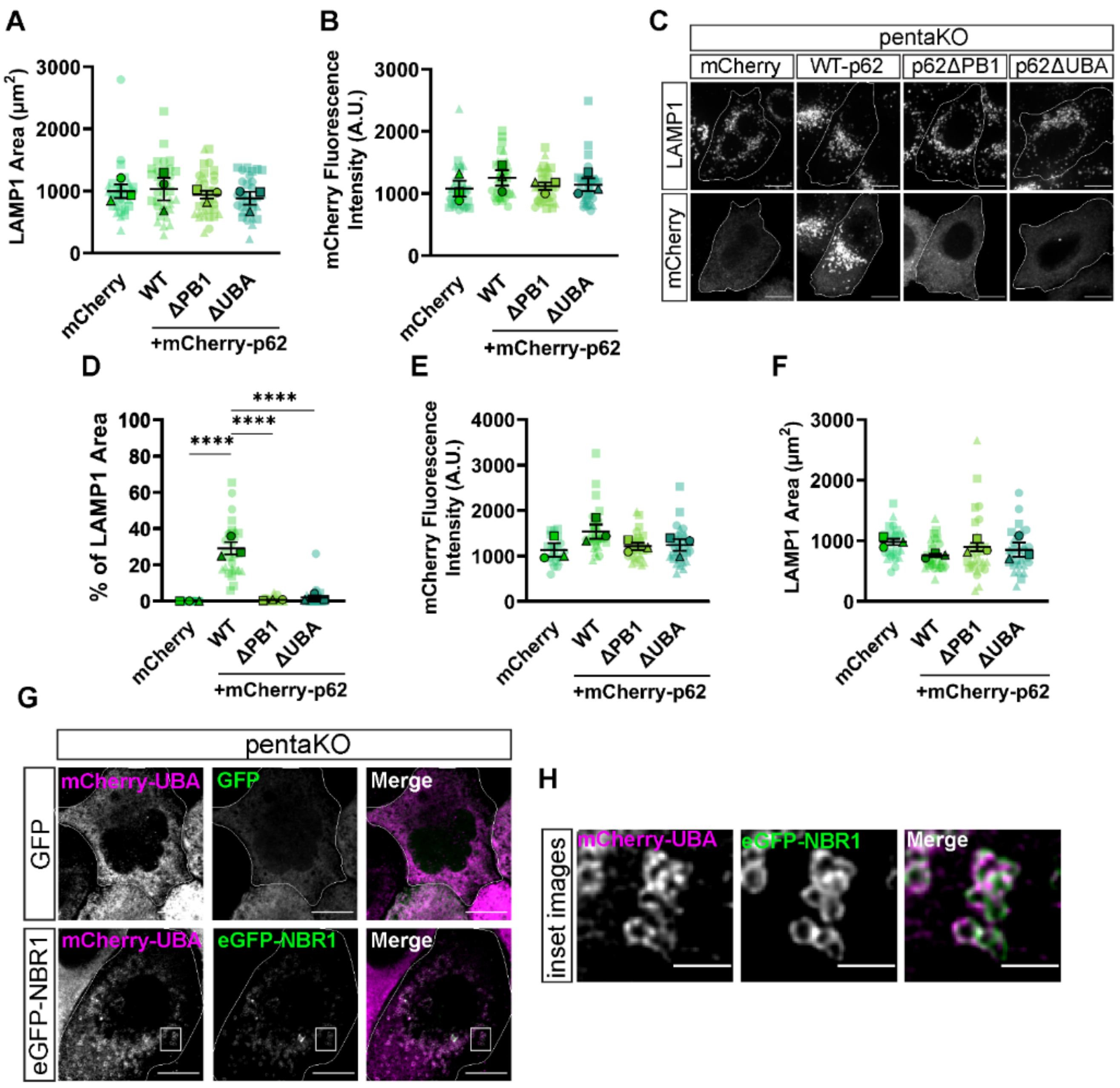


Supplemental Figure 5 – **Loss of the UBA domain decreases p62 localization to damaged lysosomes in the absence of other autophagy receptors [**[Related to Figure 4].

(A) Quantification of LAMP1 area across different conditions (One-way ANOVA, $p_{(WT/mCherry)} = 0.9941$, $p_{(WT/\Delta PB1)} = 0.9166$, $p_{(WT/\Delta UBA)} = 0.7421$, N=3, $n_{mCherry} = 33$, $n_{WT} = 34$, $n_{\Delta PB1} = 34$, $n_{\Delta UBA} = 33$).
(B) Quantification of mCherry fluorescence intensity across different conditions (One-way ANOVA, $p_{(WT/mCherry)} = 0.5456$, $p_{(WT/\Delta PB1)} = 0.7110$, $p_{(WT/\Delta UBA)} = 0.8002$, N=3, $n_{mCherry} = 33$, $n_{WT} = 34$, $n_{\Delta PB1} = 34$, $n_{\Delta UBA} = 33$).
(C) Representative images of anti-LAMP1 and either mCherry vector WT-p62, p62ΔPB1, or p62ΔUBA in pentaKO HeLa cells. Cells were treated with LLOMe for two hours. Cell outlines are traced. Scale Bar: 10μm.
(D) Quantification of LAMP1 area overlapping with p62 area across different conditions (One-way ANOVA, $p_{(WT/mCherry)} < 0.0001$, $p_{(WT/\Delta PB1)} < 0.0001$, $p_{(WT/\Delta UBA)} < 0.0001$, N=3, $n_{mCherry} = 24$, $n_{WT} = 31$, $n_{\Delta PB1} = 29$, $n_{\Delta UBA} = 29$).
(E) Quantification of LAMP1 area across different conditions (One-way ANOVA, $p_{(WT/mCherry)} = 0.9941$, $p_{(WT/\Delta PB1)} = 0.9166$, $p_{(WT/\Delta UBA)} = 0.7421$, N=3, $n_{mCherry} = 24$, $n_{WT} = 31$, $n_{\Delta PB1} = 29$, $n_{\Delta UBA} = 29$).

(F) Quantification of mCherry fluorescence intensity across different conditions (One-way ANOVA, $p_{(WT/mCherry)}$ = 0.5456, $p_{(WT/\Delta PB1)}$ = 0.7110, $p_{(WT/\Delta UBA)}$ = 0.8002, N=3, $n_{mCherry}$ = 24, $n_{WT}$ = 31, $n_{\Delta PB1}$ = 29, $n_{\Delta UBA}$ = 29).
(G) Representative images of pentaKO HeLa expressing p62ΔUBA and either GFP or eGFP-NBR1. Cell outlines are traced. White box indicates inset region. Scale Bar: 10μm.
(H) Inset images of (G). Scale Bar: 2μm.

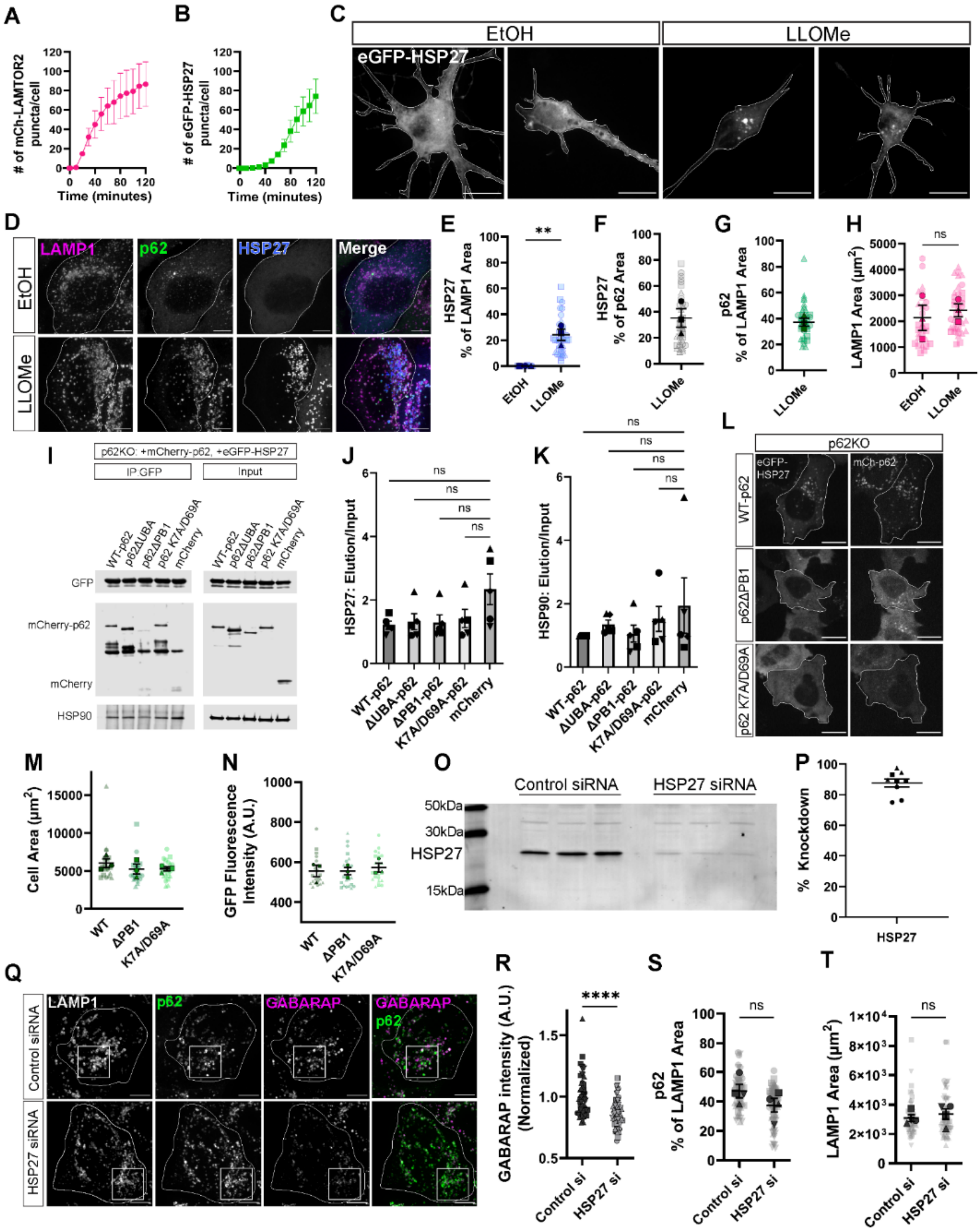
A
### of mCh-LAMTOR2 puncta/cell
Time (minutes)
B
### of eGFP-HSP27 puncta/cell
Time (minutes)
C
EtOH
LLOMe
eGFP-HSP27
D
LAMP1
p62
HSP27
Merge
EtOH
LLOMe
E
HSP27 % of LAMP1 Area
**
F
HSP27 % of p62 Area
G
p62 % of LAMP1 Area
H
LAMP1 Area (μm2)
ns
I
p62KO: +mCherry-p62, +eGFP-HSP27
IP:GFP
Input
WT-p62
p62ΔUBA
p62ΔPB1
p62 K7A/D69A
mCherry
GFP
mCherry-p62
mCherry
HSP90
J
HSP27: Elution/Input
ns
WT-p62
ΔUBA-p62
ΔPB1-p62
K7A/D69A-p62
mCherry
K
HSP90: Elution/Input
L
p62KO
eGFP-HSP27
mCh-p62
WT-p62
p62ΔPB1
p62 K7A/D69A
M
Cell Area (μm2)
WT
ΔPB1
K7A/D69A
N
GFP Fluorescence Intensity (A.U.)
O
Control siRNA
HSP27 siRNA
50kDa
30kDa
HSP27
15kDa
P
% Knockdown
HSP27
Q
LAMP1
p62
GABARAP
Control siRNA
HSP27 siRNA
R
GABARAP intensity (A.U.) (Normalized)
****
Control si
HSP27 si
S
p62 % of LAMP1 Area
T
LAMP1 Area (μm2)

Supplemental Figure 6 – **HSP27 dynamically regulates p62 in lysophagy** [Related to Figure 6].
(A) Quantification of total mCherry-LAMTOR2 puncta per cell in HeLa cells following treatment with 750μM LLOMe (N=7).
(B) Quantification of total eGFP-HSP27 puncta per cell in HeLa cells following treatment with 750μM LLOMe (N=7).
(C) Representative images of $i^3$Neurons expressing eGFP-HSP27 following treatment with either EtOH or 1.0mM LLOMe for two hours. Cell outlines are traced. Scale Bar: 10μm.
(D) Representative images of HeLa expressing eGFP-HSP27 co-localizing with anti-LAMP1 and anti-p62 following treatment with EtOH or 750μM LLOMe. Cells were treated for two hours. Cell outlines are traced. Scale Bar: 10μm.
(E) Quantification of eGFP-HSP27 occupying anti-LAMP1 area across different conditions (Unpaired t-test, $p < 0.01$, N=3, $n_{EtOH} = 29$, $n_{LLOMe} = 39$).
(F) Quantification of eGFP-HSP27 occupying anti-p62 area in HeLa cells treated with LLOMe (HSP27: 35 ± 7.2% of p62 area).
(G) Quantification of anti-p62 occupying anti-LAMP1 area in HeLa cells treated with LLOMe (p62: 37 ± 3.0% of LAMP1 area).
(H) Quantification of anti-LAMP1 area in HeLa-M cells treated with either EtOH or 750μM LLOMe (Unpaired t-test, $p = 0.6242$, N = 3, $n_{EtOH} = 29$, $n_{LLOMe} = 39$).
(I) Uncropped representative western blot demonstrating eGFP-HSP27 co-immunoprecipitation of either WT, p62ΔUBA, p62ΔPB1, K7A/D69A p62, or mCherry vector.
(J) Quantification of immunoprecipitation of eGFP-HSP27 across different conditions (Kruskal-Wallis test, $p_{(mCherry/WT\text{-}p62)} = 0.2347$, $p_{(mCherry/\Delta UBA)} = 0.1924$, $p_{(mCherry/\Delta PB1)} = 0.1924$, $p_{(mCherry/(K7A/D69A))} = 0.5762$, N=5).
(K) Quantification of co-immunoprecipitation of endogenous HSP90 with eGFP-HSP27 (Kruskal-Wallis test, $p_{(mCherry/WT\text{-}p62)} > 0.9999$, $p_{(mCherry/\Delta UBA)} > 0.9999$, $p_{(mCherry/\Delta PB1)} > 0.9999$, $p_{(mCherry/(K7A/D69A))} > 0.9999$, N=5).
(L) Representative images of eGFP-HSP27 and mCherry-p62 in p62KO HeLa cells. Cells were transfected with either WT, p62ΔPB1, or K7A/D69A p62. Cells were treated with 750μM LLOMe for two hours. Cell outlines are traced. Scale Bar: 10μm.
(M) Quantification of the cell area per cell across conditions (One-way ANOVA, $p_{(WT/\Delta PB1)} = 0.5725$, $p_{(WT/(K7A/D69A))} = 0.5899$, $p_{(\Delta PB1/(K7A/D69A)} = 0.9995$, N=3, $n_{WT} = 21$, $n_{\Delta PB1} = 22$, $n_{K7A/D69A} = 23$).
(N) Quantification of GFP fluorescence intensity per cell across conditions (One-way ANOVA, $p_{(WT/\Delta PB1)} = 0.9997$, $p_{(WT/(K7A/D69A))} = 0.8625$, $p_{(\Delta PB1/(K7A/D69A)} = 0.8506$, N=3, $n_{WT} = 21$, $n_{\Delta PB1} = 22$, $n_{K7A/D69A} = 23$).
(O) Western blot analysis of cell lysates from HeLa cells, evaluating expression of HSP27 following transient transfection of either control siRNA or HSP27 pooled siRNA.
(P) Quantification of percentage knockdown in HeLa-M cells that were transfected with Hspb1 pooled siRNA relative to total protein and to control siRNA (88 ± 2.6% knock down).
(Q) Representative images of anti-p62 and anti-GABARAP in HeLa cells following transient transfection of control siRNA or Hspb1 siRNA. White box indicates inset region. Scale bar: 10μm.
(R) Quantification of anti-GABARAP fluorescence intensity at anti-p62 segmented area across conditions (Mann-Whitney test, $p < 0.0001$, $N_{Control} = 57$, $N_{Hspb1} = 71$).
(S) Quantification of anti-p62 area overlapping anti-LAMP1 area across conditions (Unpaired t-test, $p = 0.1877$, N = 4, $n_{Control} = 57$, $n_{Hspb1} = 71$).
(T) Quantification of LAMP1 area per cell across conditions (Unpaired t-test, $p = 0.5532$, N = 4, $n_{Control} = 57$, $n_{Hspb1} = 71$).

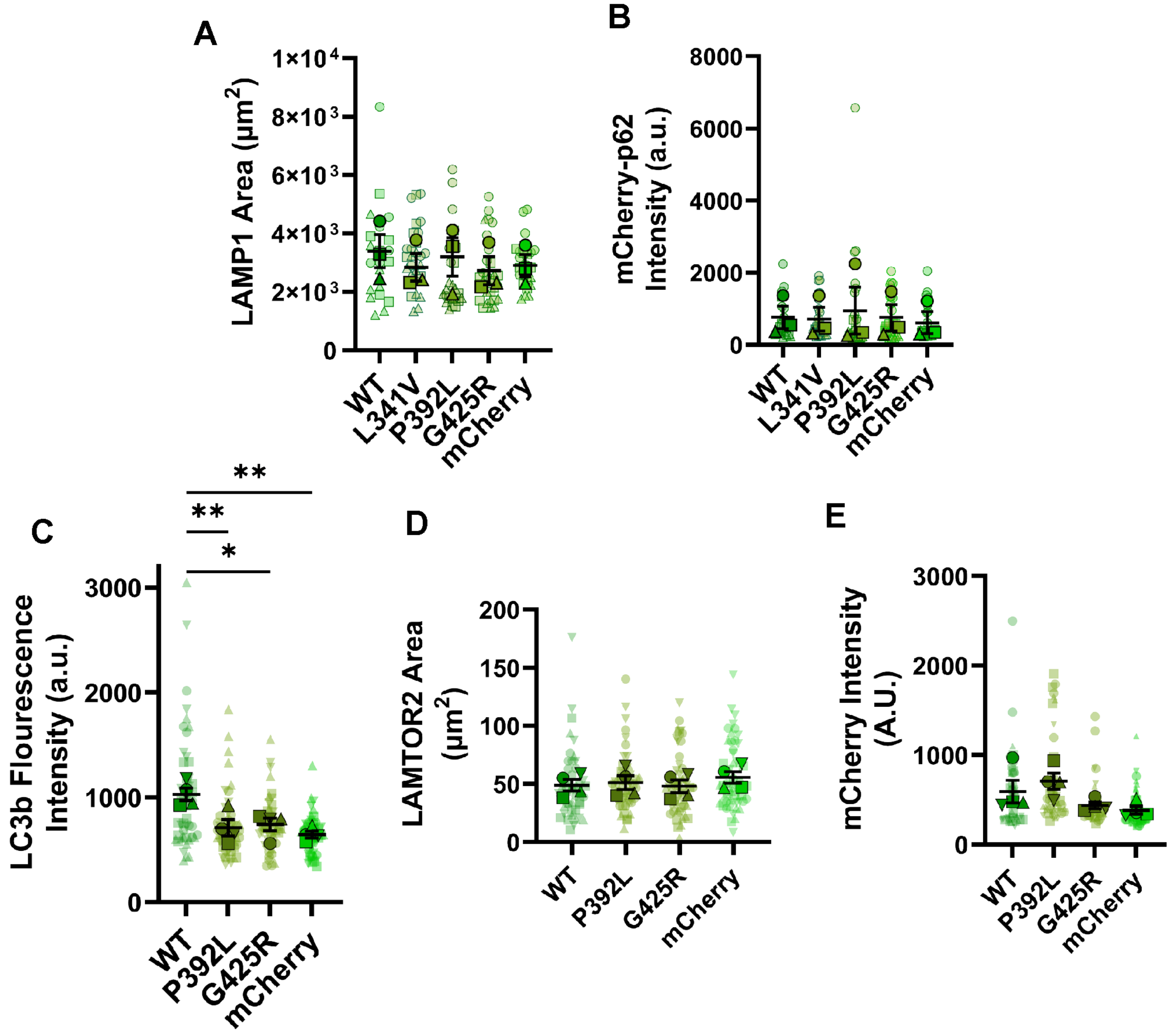


Supplemental Figure 7 – **ALS-associated mutations in p62 disrupt lysophagy** [Related to Figure 7].
(A) Quantification of LAMP1 area across different conditions (One-way ANOVA, $p_{(WT/L341V)}$ = 0.8636, $p_{(WT/P392L)}$ = 0.9963, $p_{(WT/G425R)}$ = 0.7764, $p_{(WT/mCherry)}$ = 0.9040, N=3, $n_{WT}$ = 22, $n_{L341V}$ = 27, $n_{P392L}$ = 27, $n_{G425R}$ = 27, $n_{mCherry}$ =25).
(B) Quantification of mCherry fluorescence intensity across different conditions (One-way ANOVA, $p_{(WT/L341V)}$ = 0.9541, $p_{(WT/P392L)}$ = 0.9883, $p_{(WT/G425R)}$ = 0.9791, $p_{(WT/mCherry)}$ = 0.5829, N=3, $n_{WT}$ = 22, $n_{L341V}$ = 27, $n_{P392L}$ = 27, $n_{G425R}$ = 27, $n_{mCherry}$ =25).
(C) Quantification of LC3b intensity at LAMTOR2 puncta across different conditions (One-way ANOVA, $p_{(WT/P392L)}$ < 0.01, $p_{(WT/G425R)}$ < 0.05, $p_{(WT/mCherry)}$ < 0.01, N=4, $n_{WT}$ = 49, $n_{P392L}$ = 51, $n_{G425R}$ = 51, $n_{mCherry}$ = 49).
(D) Quantification of LAMTOR2 area across different conditions (One-way ANOVA, $p_{(WT/P392L)}$ = 0.9849, $p_{(WT/G425R)}$ = 0.9983, $p_{(WT/mCherry)}$ 0.7200, N=4, $n_{WT}$ = 49, $n_{P392L}$ = 51, $n_{G425R}$ = 51, $n_{mCherry}$ = 49).
(E) Quantification of mCherry fluorescence intensity across different conditions (One-way ANOVA, $p_{(WT/P392L)}$ = 0.6312, $p_{(WT/G425R)}$ = 0.4536, $p_{(WT/mCherry)}$ = 0.2341, N=4, $n_{WT}$ = 49, $n_{P392L}$ = 51, $n_{G425R}$ = 51, $n_{mCherry}$ = 49).
